# β-importins Tnpo-SR and Cadmus and the small GTPase Ran promote ovarian cyst formation in *Drosophila*

**DOI:** 10.1101/2021.01.31.429051

**Authors:** Allison N. Beachum, Taylor D. Hinnant, Anna E. Williams, Amanda M. Powell, Elizabeth T. Ables

**Affiliations:** Department of Biology, East Carolina University, Greenville, NC 27858

**Keywords:** Oocyte, oogenesis, germline, cyst, germline stem cell

## Abstract

Germ cells undergo mitotic expansion via incomplete cytokinesis, forming cysts of undifferentiated cells that remain interconnected prior to meiotic initiation, through mechanisms that are not well-defined. In somatic cells, Ras-related nuclear protein (Ran) spatiotemporally regulates mitotic spindle assembly, cleavage furrow formation and abscission. Here, we identify Ran and β-importins as critical regulators of cyst development in the *Drosophila* ovary. Depletion of *Ran* or the β-importins *Tnpo-SR* and *cadmus* disrupts oocyte selection and results in egg chambers with variable numbers of germ cells, suggesting abnormal cyst development and cyst fragmentation. We demonstrate that Ran, Tnpo-SR, and Cadmus regulate key cellular processes during cyst formation, including cell cycle dynamics, fusome biogenesis, and ring canal stability, yet do so independently of mitotic spindle assembly. Further, Tnpo-SR and Cadmus control cyclin accumulation and suppress cytokinesis independent of Ran-GTP, suggesting that β-importins sequester protein cargos that normally promote the mitotic-to-meiotic transition. Our data demonstrates that Ran and β-importins are critical for germ cell cyst formation, a role that is likely conserved in other organisms.

**SUMMARY STATEMENT:** Ran and two β-importins function coordinately to promote oocyte selection and cyst development in the *Drosophila* ovary.

## INTRODUCTION

Germ cells develop as cysts of interconnected undifferentiated cells (Lu et al., 2017; Matova and Cooley, 2001; Pepling and Lei, 2018). Cysts arise via incomplete cytokinesis, manifested by cleavage furrow arrest and the formation of stable ring canals that permit transport of cytoplasm and organelles (de Cuevas et al., 1997; Guertin et al., 2002; Haglund et al., 2011; Pepling and Lei, 2018). Cyst formation increases cytoplasmic volume, enhances sensitivity to DNA damage, and ensures robust oocyte development (Chaigne et al., 2020; de Cuevas et al., 1997; Pepling and Lei, 2018; Pepling et al., 1999; Yamashita, 2018). Cyst formation is evolutionarily conserved and essential for optimal fertility in multiple phyla (Haglund et al., 2011; Lu et al., 2017).

The fruit fly, *Drosophila melanogaster*, is an elegant model of germline cyst formation. In adult females, oocytes are produced in a linear spatiotemporal arrangement fueled by the activity of germline stem cells (GSCs; Figure 1A) (Hinnant et al., 2020). GSCs divide asymmetrically, giving rise to one daughter that remains a GSC and a cystoblast that differentiates. Delayed abscission between the GSC and cystoblast keeps the pair connected well into G2 of the subsequent cell cycle (Ables and Drummond-Barbosa, 2013; Eikenes Å et al., 2015; Hinnant et al., 2017; Mathieu et al., 2013; Matias et al., 2015). Cystoblasts divide four times with incomplete cytokinesis, creating 16-cell cysts. At the completion of the mitotic divisions, cleavage furrows between germ cells are modified into stable ring canals (Grieder et al., 2000; Guertin et al., 2002; Hinnant et al., 2020). The fusome, a cytoplasmic organelle composed of microtubules, endoplasmic reticulum-derived vesicles, and membrane skeleton proteins, traverses the intercellular bridges (Hinnant et al., 2020; Ong and Tan, 2010; Pepling and Lei, 2018; Röper, 2007; Snapp et al., 2004). The fusome branches as mitotic divisions progress, providing a synchronization hub for cell cycle regulation and cell polarity (Grieder et al., 2000; Hinnant et al., 2020; Lighthouse et al., 2008). These specialized cytological features of germ cells facilitate interconnectivity and are vital to oocyte selection; however, the molecular mechanisms underlying fusome morphogenesis and incomplete cytokinesis are not well understood.

**Figure 1.**
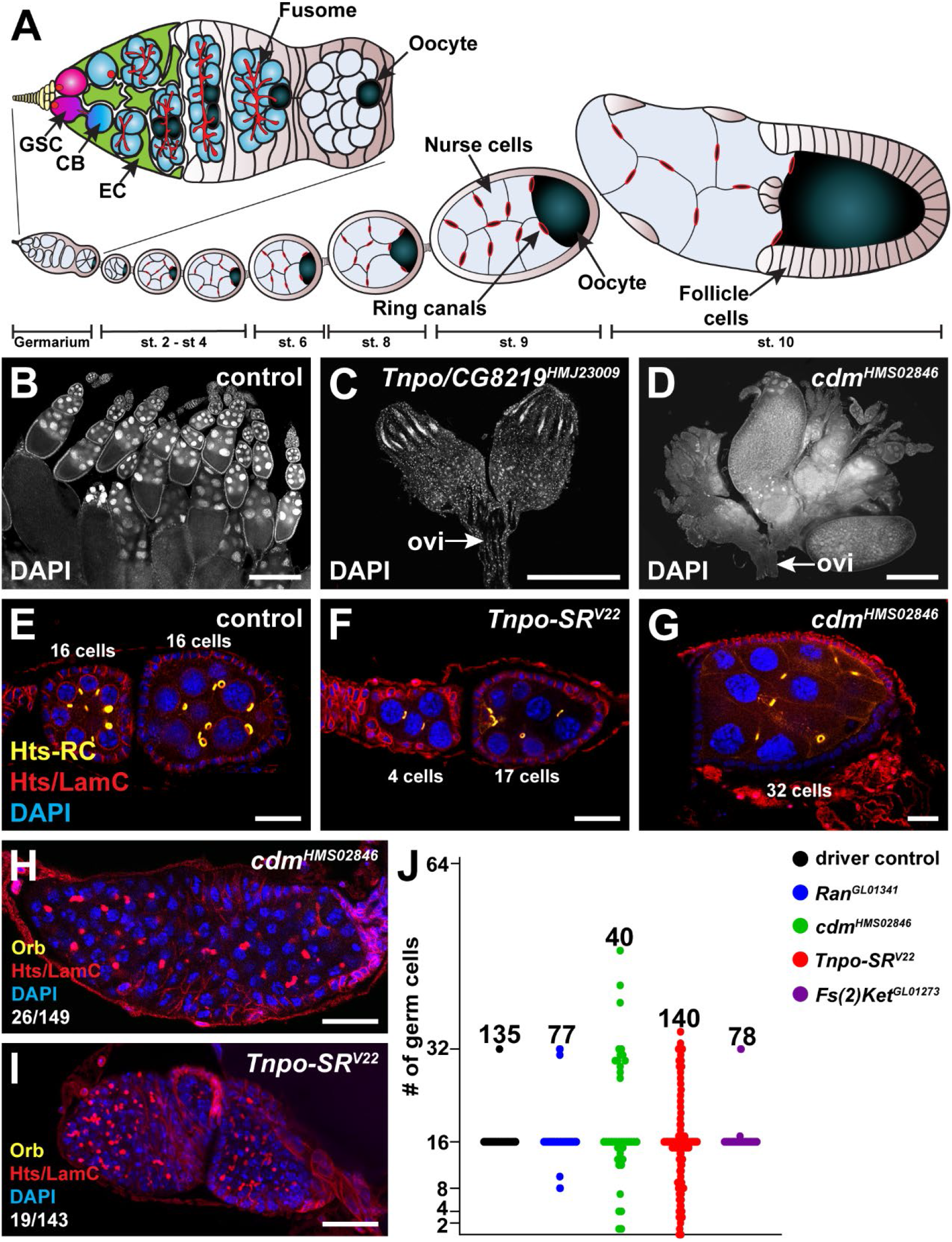
An RNAi screen identified Ran and β-importins necessary for germ cell development. (A) The *Drosophila* ovary is composed of ovarioles harboring a germarium (top) and older egg chambers containing developing oocytes (bottom). In the germarium, germline stem cells (GSCs; pink) are adjacent to cap cells (yellow) and escort cells (green). GSCs divide to form cystoblasts (blue) that divide four times to form 16-cell cysts (blue). Cysts are composed of 15 nurse cells and one oocyte (dark blue), connected by 15 intercellular bridges that can be visualized by ring canals and the fusome (red) that transects each bridge. (B-D) Maximum intensity projections of *nos-Gal4* control (B), *nos-Gal4>Tnpo/CG82l9^HMJ23009^* (C), or *nos-Gal4>cdm^HMS02846^* (D) knockdown ovaries labeled with DAPI (white; nuclei). Arrows indicate oviducts (ovi). Scale bar, 200 μm. (E-G) Egg chambers of driver control (E), *nos-Gal4>Tnpo-SR^V22^* (F), and *nos-Gal4>cdm™^HMS02846^* (G) ovarioles labeled with anti-Hts-RC (yellow), anti-Hts (red), anti-LamC (red), and DAPI (blue). (H-I) *nos-Gal4>Tnpo-SR^V22^* (H) and *nos-Gal4>cdm^HMS02846^* (I) females labeled with anti-Orb (yellow; oocyte), anti-Hts and anti-LamC (red), and DAPI (blue). Values in bottom left corner of G-H represent the number of ovarioles containing germline tumors over the total number of ovarioles analyzed. Scale bar, 20 μm. (J) Number of germ cells per egg chamber at 6 days after eclosion. Each dot represents one egg chamber; numbers above dots represent the total number of egg chambers analyzed. Quantifications do not include tumorous egg chambers.

Formation of an actomyosin-based contractile network at the equatorial cleavage plane during cytokinesis is spatiotemporally regulated and requires a dynamic interplay between the cytoskeleton, the mitotic microtubule-based central spindle, and membrane trafficking (Frémont and Echard, 2018; Haglund et al., 2011; Nguyen and Robinson, 2020; Pollard and O’Shaughnessy, 2019). Central to this process is Ras-related nuclear protein (Ran), a small guanosine triphosphate (GTP)-binding protein implicated in protein transport during mitosis and interphase (Ozugergin and Piekny, 2020; Sazer and Dasso, 2000). Ran partners with α- and β-importins to move protein cargo into the nucleus and promote mitotic spindle assembly (Cautain et al., 2015; Okada et al., 2008). Ran also regulates cell polarization and contractile protein localization during cleavage furrow ingression (Beaudet et al., 2017; Silverman-Gavrila et al., 2008). Chromatin-associated Ran-GTP may help recruit and organize essential proteins at the cortex, thereby ensuring the robustness of cytokinesis (Nguyen and Robinson, 2020).

Here, we identify Ran and β-importins as novel regulators of germline cyst formation in the *Drosophila* ovary. We show that Ran and β-importins Tnpo-SR and Cadmus (Cdm) control cyst formation by regulating cell cycle dynamics, fusome biogenesis, and ring canal stability. Ran function depends on its active, GTP-bound state, as expression of a dominant-negative *Ran* mutant recapitulates *Ran* loss-of-function. *Ran*, *Tnpo-SR*, and *cdm* are not essential for mitotic spindle assembly in undifferentiated germ cells. Moreover, knock-down of *Tnpo-SR* or *cdm* does not completely phenocopy loss of *Ran*, suggesting that β-importins control cell cycle progression and suppress cytokinesis independent of Ran activity. Given the conservation of germline cyst formation across phyla, our data suggest that Ran and specific β-importins may play similar roles in other organisms.

## RESULTS

### RNAi screen reveals that β-importins and Ran are essential for germ cell development

We previously identified *Transportin-Serine/Arginine rich (Tnpo-SR)* in a genetic mosaic screen for genes controlling GSC self-renewal (Ables et al., 2016). *Tnpo-SR*, homologous to mammalian *TNPO3*, encodes a β-importin (Allemand et al., 2002; Lai et al., 2001). To determine whether *Tnpo-SR* loss-of-function phenotypes reflect a general role for importins in germ cell mitoses, we performed a secondary reverse genetic screen using germline-enhanced RNA interference (RNAi) to knock-down expression of nine *Drosophila* importins and the small GTPase *Ran* specifically in germ cells (Figure 1B-D, Table 1) (Ni et al., 2011; Rørth, 1998; Van Doren et al., 1998). In contrast to driver-only controls (Figure 1B, Table 1), three RNAi lines produced completely agametic ovaries, defined by the absence of egg chambers (Figure 1C). More intriguingly, four RNAi lines corresponding to *Ran* and the β-importins *Tnpo-SR*, *cadmus* (*cdm*), and *Fs(2)Ket* (also called *Ketel)* exhibited a partial agametic phenotype, in which germ cells were present in some ovarioles, but in unusually shaped egg chambers (Figure 1D; Table 1). Cdm and Tnpo-SR are paralogs, sharing 42% amino acid similarity, while Ketel is less conserved. These results demonstrate that Ran and some β-importins are necessary for proper oocyte development and may have distinct functions.

**Table 1.** Frequency of ovarioles or egg chambers that showcase germline phenotypes in β-importin- and Ran-depleted ovaries.

| genotype | % of agametic ovarioles | average # of GSCs at 3 days after eclosion<sup>a</sup> | % of egg chambers that do not equal 16 cells | % of egg chambers that do not equal 2<sup>n</sup> germ cells | % of 16-cell egg chambers without an oocyte |
| --- | --- | --- | --- | --- | --- |
| driver control | 0 | 2.8 | 0.64 | 0 | 0 |
| *Fs(2)Ket<sup>GL01273</sup>* | 3.6 | 1.8 *** | 3.9 | 1.3 | 0 |
| *Ran<sup>GL01341</sup>* | 1.5 | 1.1 *** | 11.31*** | 2.6 | 4.0 |
| *cdm<sup>HMS02846</sup>* | 33.1 | 0.27 *** | 32.0 *** | 55.0 | 0 |
| *Tnpo-SR<sup>V22</sup>* | 2.5 | 1.5 *** | 46.0 *** | 29.3 | 33.8 |
| *Ran<sup>HA.1</sup>* | 0 | 2.9 | 0 | 0 | 0 |
| *Ran<sup>HA.2</sup>* | 0 | 2.4 | 0 | 0 | 0 |
| *Ran<sup>T24N.1</sup>* | 63.8 | 0.25 *** | 15.6 *** | 12.5 | 23.5 |
| *Ran<sup>T24N.2</sup>* | 49.6 | 0.38 *** | 19.5*** | 9.7 | 11.5 |
| *Ran<sup>Q69L.1</sup>* | 0 | 1.9 *** | 0 | 0 | 0 |
| *Ran<sup>Q69L.2</sup>* | 2.0 | 2.0 *** | 6.4 ** | 2.2 | 0 |

<sup>a</sup>See Suppl. Figure S1.
\*\* $p < 0.01$ , \*\*\* $p < 0.001$ . Students two-tailed t-test and Chi-squared test.

Since GSCs give rise to all germ cells, we hypothesized that agametic ovaries resulting from *Ran* and *β-importin* knock-down could arise from a failure to establish or maintain GSCs. To test this, we quantified the number of GSCs per germarium. GSCs and their progeny can be easily identified by immunolabeling of Hts, a fusome-resident protein in germ cells, and LaminC (LamC), a nuclear envelope protein highly expressed in cap cells (the major cellular component of the GSC niche) (Ables and Drummond-Barbosa, 2013). Unlike cytoblasts, GSCs possess an anteriorly localized fusome juxtaposed to the cap cells (de Cuevas and Spradling, 1998; Lin et al., 1994). All four partially agametic RNAi lines had significantly less GSCs per germarium at three days after eclosion than age-matched driver-only controls (Table 1; Supp. Figure 1A-C). Fewer GSCs in early adulthood suggests that *Ran*, *Tnpo-SR, Ketel*, and *cdm* are required for GSC establishment during larval/pupal development or during the proliferative surge in GSCs following the female’s first mating (Ameku and Niwa, 2016; Van Doren et al., 1998). Conditional knock-out of *Tnpo-SR* further demonstrated the requirement for β-importin function in GSC maintenance. We used two strong loss-of-function transposon insertion alleles of *Tnpo-SR (Tnpo-SR^KG04870^* and *Tnpo-SR^LL05552^)* carrying *FRT* sites to generate germline mutant mosaic germaria under the *Flippase* (*FLP*)/*FLP Recognition Target* (*FRT*) genetic mosaic lineage-tracing system (Laws and Drummond-Barbosa, 2015; Xu and Rubin, 1993). GFP-negative GSC clones were generated at equivalent rates in control, *Tnpo-SR^KG04870^*, and *Tnpo-SR^LL05552^* mosaic germaria at four days after clone induction (Supp. Figure 1F). In contrast to mock controls, *Tnpo-SR^KG04870^* and *Tnpo-SR^LL05552^* germline mutant mosaic germaria rapidly lost marked GSCs over time (Supp. Figure 1D-F), indicating that *Tnpo-SR* is cell-autonomously required for GSC self-renewal.

### Loss-of-function mutations in Ran and β-importins alter the number of germ cells per egg chamber

The intriguing phenotype resulting from knock-down of *Ran* and β-importins prompted us to examine egg chamber morphology more closely. To visualize germline and follicle cells, we combined anti-Hts antibodies with anti-Hts-RC antibodies, which are specific for a Hts cleavage product that localizes to ring canals (Petrella et al., 2007; Robinson et al., 1994). Unlike driver control egg chambers (Figure 1E; Table 1), knock-down of *Ran, cdm*, or *Tnpo-SR* significantly decreased the number of egg chambers with 16 germ cells (Figure 1F-G; Table 1). Depletion of *Ketel* did not significantly impact egg chamber formation (Table 1).

Cystoblasts normally divide synchronously four times, following a 2^n^ pattern, to produce cysts with 16 cells (King, 1970). To determine whether extra germ cells in *Ran*, *cdm*, and *Tnpo-SR* depleted ovarioles arose by alterations in the number of mitotic divisions, we quantified the number of germ cells per cyst (Figure 1J). In the rare instances where control or *Ketel* RNAi egg chambers had an abnormal number of germ cells, they invariably contained 32 cells (Figure 1J). In contrast, knock-down of *Ran*, *cdm*, or *Tnpo-SR* produced egg chambers with variable numbers of germ cells, ranging anywhere from one to 49 cells per cyst (Figure 1J). While some egg chambers housed germ cells on the order of 2^n^, a substantial number deviated from this pattern (Table 1). Germ cell numbers in adjacent chambers in RNAi-depleted egg chambers did not add up to 8, 16, or 32 cells, arguing that encapsulation defects alone are not the source of the unusual germ cell number phenotype. Some egg chambers with 32 or more cells had oocytes with five ring canals, suggesting that some abnormal egg chambers arose by an extra mitotic division. Other egg chambers, however, likely arose because of other cellular defects. Moreover, a small percentage of *cdm* and *Tnpo-SR* RNAi knock-down ovarioles (17.4% and 13.3% of all ovarioles, respectively) contained germline tumors, in which a normal follicle cell monolayer surrounded mitotically-active germ cells with underdeveloped fusomes (Figure 1H-I; Supp. Figure 2A-C’). In *Tnpo-SR* RNAi ovarioles, undifferentiated germ cells were encapsulated with more-developed germ cells (Supp. Figure 2A). Germline tumors contained mitotically-active cells with short or unbranched fusomes and immature ring canals that frequently appeared as dot-like structures, reminiscent of the abscission site in GSC/cystoblast pairs (Supp. Figure 2B-C’). These structures suggest tumors arose because of improper abscission events in 2-cell and/or 4-cell cysts. Immunostaining for Sex-lethal (Sxl), which is enriched in GSCs and cystoblasts in the anterior-most region of the germarium (Supp. Figure 2D-D’) (Chau et al., 2009), confirmed that *Tnpo-SR* RNAi germ cells have not completed differentiation (Supp. Figure 2E-E’). We conclude that Ran, Tnpo-SR, and Cdm impact the synchrony and number of cyst divisions.

**Figure 2.**
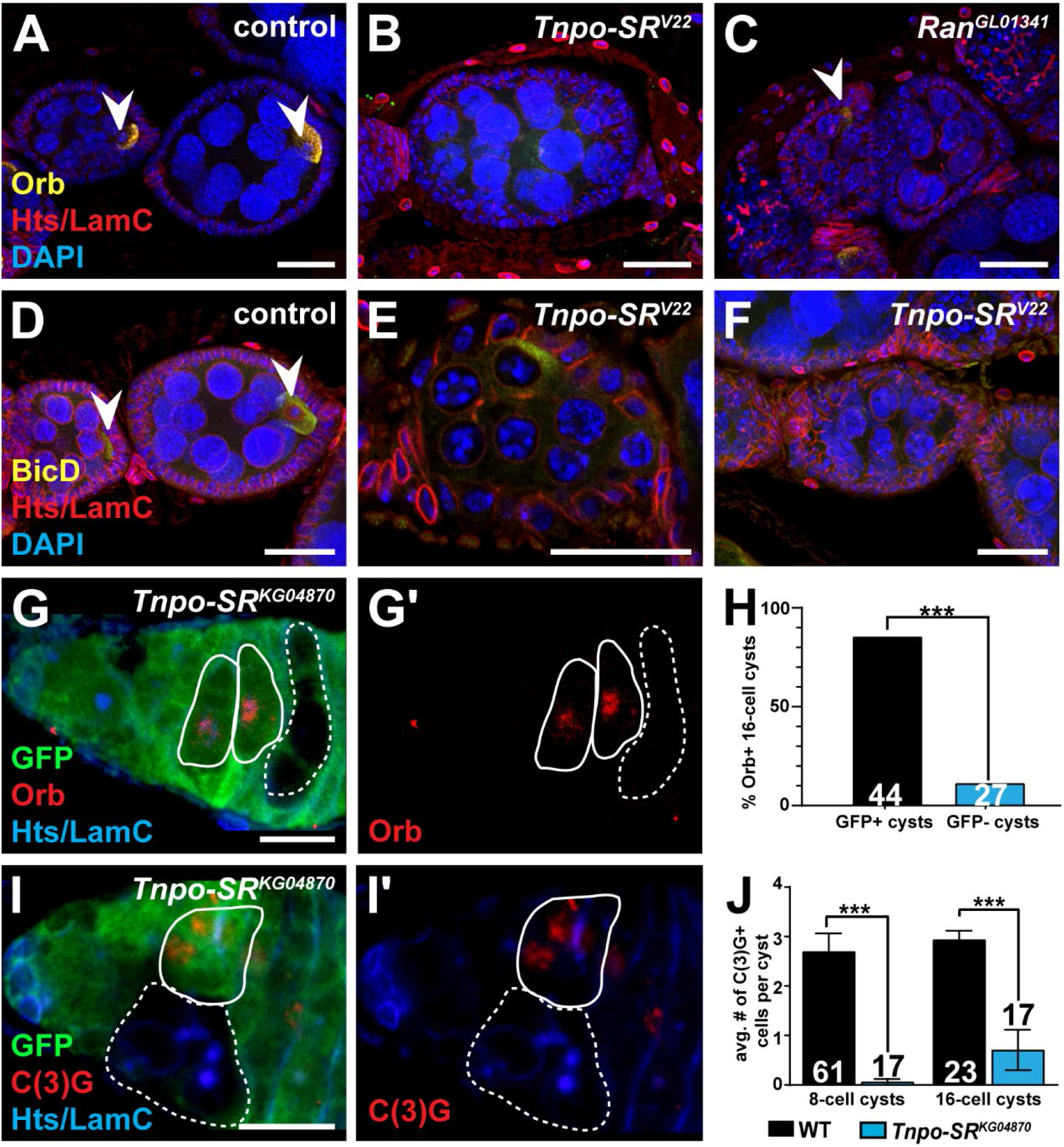
*Ran* and *Tnpo-SR* are required for proper oocyte selection. (A-C) Driver control (A), *nos-Gal4>Tnpo-SR^V22^* (B), and *nos-Gαl4>Rαn^GL01341^* egg chambers labeled with anti-Orb (yellow), anti-Hts (red), anti-LamC (red), and DAPI (blue). (D-F) Driver control (D) and *nos-Gal4>Tnpo-SR^V22^* (E-F) egg chambers labeled with anti-BicD (yellow), anti-Hts (red), anti-LamC (red), and DAPI (blue). Arrowheads indicate oocytes, recognized by their condensed nuclei. (G) Frequencies of egg chambers containing 0 (grey), 1 (white), or 2 (black) oocytes. The numbers in the bars represent the total number of egg chambers analyzed. (H-J) *Tnpo-SR^KG04870^* mutant mosaic germarium at 8 days after clone induction labeled with anti-GFP (green; wild-type cells), anti-Hts and anti-LamC (blue), and anti-Orb (red) (H-H’) or anti-C(3)G (red). (J-J’). White lines demarcate wild-type (solid) or mutant (dashed) cysts. (I) Percentage of GFP-positive (wild-type) or GFP-negative *(Tnpo-SR^KG04870^)* 16-cell cysts with an Orb-positive oocyte 8 days after clone induction. (K) Average number of C(3)G-positive germ cells in GFP-positive (wild-type) or GFP-negative (*Tnpo-SR^KG04870^*) 8- or 16-cell cysts at 8 days after clone induction. Numbers in the bars represent the total number of cysts analyzed. Scale bar, 20 μm (A-F) and 5 μm (H-J). Error bars, mean ± SEM. \*\*\**p* < 0.001; Student’s two-tailed T-test.

### Ran, Tnpo-SR, and cdm are required for oocyte selection

During germ cell mitotic divisions, three to four cyst cells initiate meiosis by assembling the synaptonemal complex; however, only one of these cells is maintained as the oocyte (Hughes et al., 2018). Since depletion of *Ran*, *Tnpo-SR*, or *cdm* altered the number of germ cells per cyst, we hypothesized that oocyte selection was also affected. In wild-type egg chambers just posterior to the germarium, the oocyte-specific proteins Orb (Figure 2A) and Bicaudal D (BicD) (Figure 2D) had clearly accumulated in one posterior cyst cell. In contrast, egg chambers from germline-depleted *Ran*, *Tnpo-SR*, or *cdm* females did not properly express Orb *(Ran^GL01341^, cdm^HMS02846^, and Tnpo-SR^V22^*) or BicD (*Tnpo-SR^V22^*) (Figure 2B-C,E-F; Table 1). Many loss-of-function egg chambers also lacked a cell with a karyosome, a clear indicator of oocyte fate (Figure 3B-C,E-F). Importantly, most of the *nos-Gal4>Ran^GL01341^* and *nos-Gal4>cdm^HMS02846^* egg chambers with 16 germ cells contained an oocyte (Table 1), suggesting that loss of oocyte identity is a secondary effect of incorrect mitotic divisions in the absence of *Ran* and *cdm*. This was not the case, however, for *Tnpo-SR*, as 34% of *nos-Gal4>Tnpo-SR^V22^* egg chambers with 16 germ cells contained only nurse cells (Table 1). Similarly, in *Tnpo-SR* germline mutant mosaic germaria, mutant cysts located posterior to wild-type cysts (thus at a more advanced stage of development) failed to up-regulate Orb expression to similar levels as adjacent wild-type cysts (Figure 2G-H). Further, a core component of the synaptonemal complex, C(3)G, normally present as early as the 4-cell cyst stage, did not accumulate in *Tnpo-SR^KG04870^* mutant cysts to equivalent levels as adjacent wild-type cysts (Figure 2I-J). This suggests that *Tnpo-SR* is specifically required for oocyte specification independent of its role in cyst divisions.

**Figure 3.**
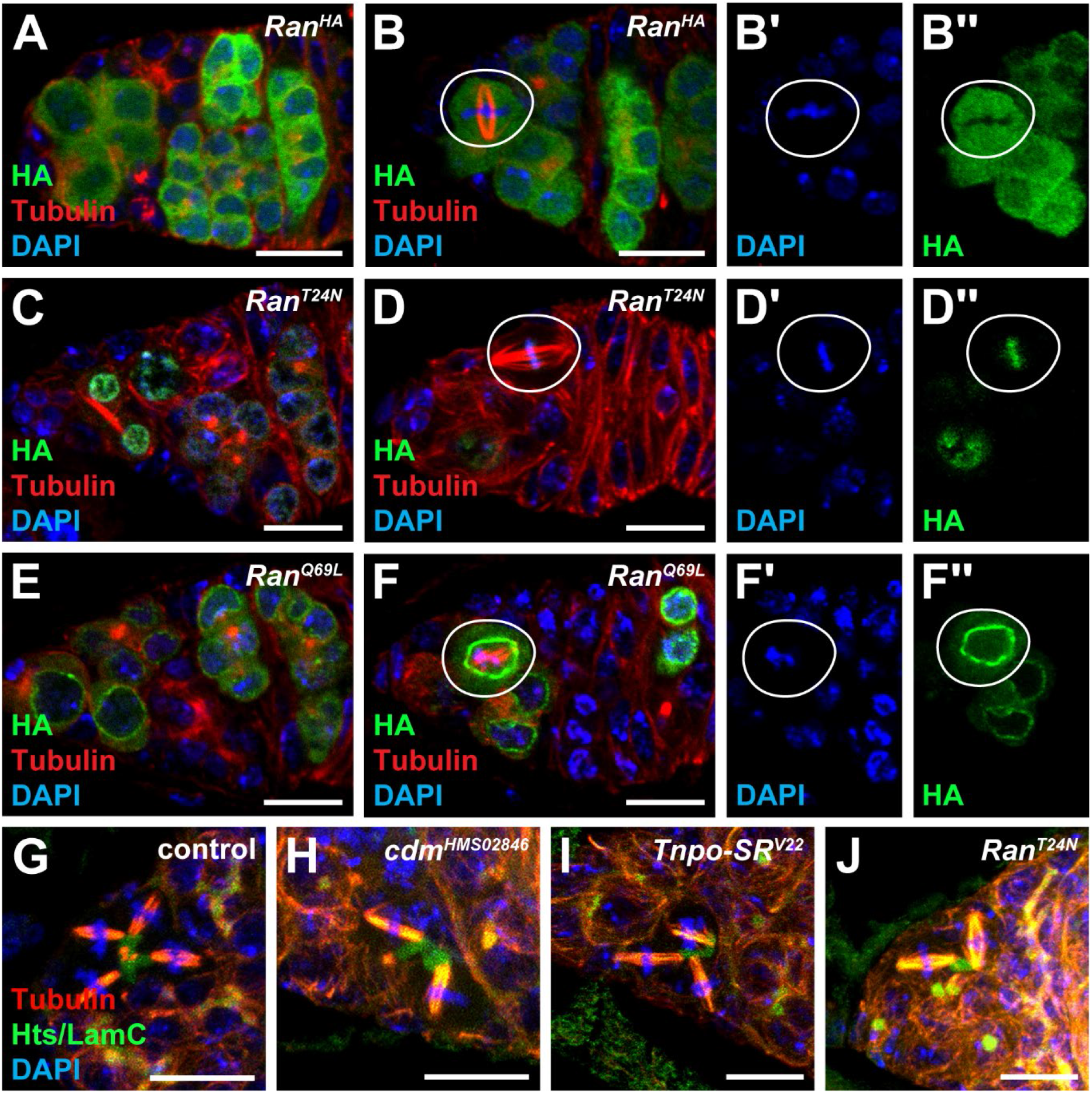
Ran is dynamically localized in germ cells but dispensable for spindle assembly. (A-F’’) Ovarioles from *nos-Gal4Ran^HA^* (A-B’’), *nos-Gal4>Ran^T24N^* (C-D’’), and *nos-Gal4>Ran^Q69L^* (E-F’’) labeled with anti-HA (green), anti-Tubulin (red) and DAPI (blue). Panels B’-B’’, D’-D’’, and F’-F’’ show the DAPI and HA channels of panels B, D, and F, respectively. (G-J) Maximum intensity projections of driver control (G), RNAi-depleted *cdm^HMS02846^* (H) and *Tnpo-SR^V22^* (I), and *Ran^T24N^* mutant (J) cysts labeled with anti-tubulin (red) anti-Hts (green), anti-LamC (green), and DAPI (blue). Scale bars, 10 μm.

### Ran is dynamically localized in cyst cells and dispensable for spindle assembly

One well-characterized function of Ran and β-importins is the regulation of spindle assembly (Cesario and McKim, 2011; Forbes et al., 2015; Nguyen and Robinson, 2020; Ozugergin and Piekny, 2020). Spindle orientation is critical for GSC maintenance, and mitotic spindle remnants accumulate in fusomes as cysts divide (Cheng et al., 2008; Deng and Lin, 1997; Grieder et al., 2000; Máthé et al., 2003). Like other small GTP-binding proteins, Ran cycles between active (GTP-bound) and inactive (GDP-bound) states (Ozugergin and Piekny, 2020). Ran-GTP dissociates importin-bound cargo in the nucleus and is converted to Ran-GDP in the cytoplasm. In some cell types, this establishes a gradient of active Ran in the cell that promotes spindle formation.

To determine whether such a gradient may exist in cyst cells, we first analyzed the subcellular distribution of Ran. We took advantage of a previously characterized transgene in which the germline-specific *UASp* promoter drives an N-terminal hemagglutinin (HA)-tagged Ran *(Ran^HA^)* (Cesario and McKim, 2011). Using *nos-Gal4* to drive *Ran^HA^* in germ cells, we then localized Ran using anti-HA antibodies in comparison to mitotic spindles (labeled with α-tubulin), fusomes, and chromosomes (Figure 4A-A’). Unlike neuroblasts, in which Ran-HA is nuclearly localized during interphase and overlaps with the mitotic spindle during mitosis (Cesario and McKim, 2011), we did not observe strong nuclear localization of Ran-HA in GSCs or cyst cells (Figure 3A-B’’). Ran-HA was higher in the cytoplasm than in nuclei of interphase and mitotically-active cells and, consistent with neuroblasts and embryonic cells (Cesario and McKim, 2011; Trieselmann and Wilde, 2002), was absent from chromosomes (Figure 3B-B’’). Ran-HA expression was higher in some cysts than others, likely due to minor variations in driver strength. In contrast to meiotic spindles (Cesario and McKim, 2011), we did not observe strong accumulation of Ran-HA surrounding mitotic spindles (Figure 3B-B’’).

**Figure 4.**
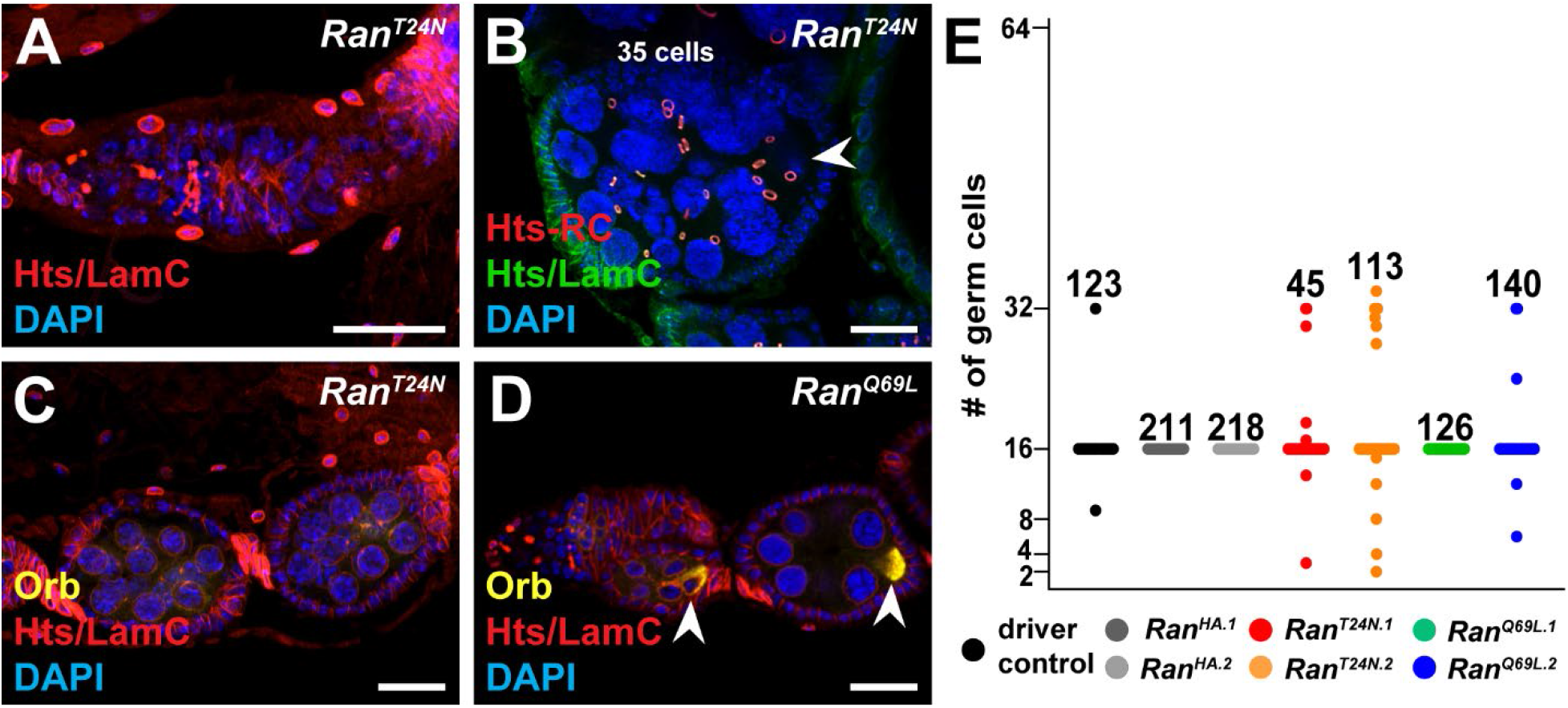
Ran function in dividing germ cells requires Ran-GTP activity. (A) Maximum intensity projection of a germarium from a *nos-Gal4>Ran^T24N^* ovary immunostained with anti-Hts (red), anti-LamC (red), and DAPI (blue). (B) Maximum intensity projection of a *Ran^T24N^* germarium stained with anti-Hts-RC (red), anti-Hts (green), anti-LamC (green), and DAPI (blue). (C-D) *Ran^T24N^* (C, GDP-locked mutant) and *Ran^Q69L^* (D, GTP-locked mutant) germaria labeled with anti-Orb (yellow), anti-Hts (red), anti-LamC (red), and DAPI (blue). Arrowheads indicate oocytes, as recognized by their condensed nuclei. Scale bars, 20 μm. (E) Number of germ cells per egg chamber at 6 days after eclosion. Each dot represents one egg chamber; numbers above dots represent the total number of egg chambers analyzed.

We then asked whether Ran-GDP accumulated in specific subcellular locations in cyst cells. We over-expressed an HA-tagged *Ran^T24N^*, which carries an amino acid substitution that locks Ran into a GDP-bound state and inhibits cargo loading (Cesario and McKim, 2011; Drutovic et al., 2020). As expected, Ran-GDP was primarily cytoplasmic in interphase cells (Figure 3C). We also observed that Ran-GDP accumulated in the nucleus of mitotically-dividing cells (Figure 3C), overlapping with chromatin at metaphase (Figure 3D-D’’). Mitotic spindles appeared to assemble correctly (see below), but chromatin was dispersed in interphase cells.

The subcellular localization of Ran-GTP was markedly different than that of Ran-GDP. We used a second HA-tagged transgene, *Ran^Q69L^*, to over-express a constitutively active form of Ran locked in a hydrolysis-resistant GTP-bound state (Cesario and McKim, 2011). Ran-GTP was excluded from cyst cell nuclei during interphase (Figure 3C) and accumulated at the presumptive nuclear membrane during mitosis (Figure 3D-D’’). Taken together, these experiments suggest that Ran aids in chromatin condensation or cytokinesis following the completion of mitosis.

Finally, we asked whether Ran and β-importins are necessary for spindle formation. Using immunostaining for α-Tubulin, we confirmed that mitotic spindles in dividing cysts are anchored to the fusome, oriented perpendicularly (Figure 3G) (Megraw and Kaufman, 2000; Storto and King, 1989). Surprisingly, in *cdm*-depleted (Figure 3H) or *Tnpo-SR-depleted* (Figure 3I) germ cells, similar-sized spindles formed and were appropriately anchored and oriented with respect to fusomes. Similarly, we were unable to find defects in spindle formation or orientation in *nosGal4>Ran^T24N^* germ cells (Figure 3D,J). We conclude that mitotic spindles can form even when Ran-GTP is depleted, suggesting that low levels of active Ran are sufficient to maintain germ cell mitoses.

### GSC establishment, cyst formation, and oocyte selection require active Ran-GTP

Given the intriguing localization of Ran-GTP, we then asked whether active Ran-GTP is required for GSC establishment and cyst formation. Flies expressing the *Ran^T24N^* and *Ran^Q69L^* mutations in germ cells were compared to flies expressing wild-type *Ran^HA^*. We expected that constitutively expressing Ran-GDP would titrate out the amount of active Ran, thus mimicking *Ran* loss-of-function. Indeed, over-expression of *Ran^T24N^* resulted in a partial agametic phenotype and the average number of GSCs per germarium was substantially reduced as compared to age-matched controls (Figure 4A; Table 1). Over-expression of dominant negative *Ran^T24N^* also altered the number of germ cells per cyst (Figure 4B-C,E) and, similar to knock-down of *Ran* via RNAi, the number of germ cells did not always follow a 2^n^ pattern (Figure 4E, Table 1). Over-expression of *Ran^T24N^* also impaired oocyte selection, as 12-24% of egg chambers with 16 germ cells lacked an Orb-positive cell (Table 1). These phenotypes are consistent with depletion of *Ran* via RNAi and likely contribute to the female sterility phenotype noted in prior studies (Cesario and McKim, 2011).

Intriguingly, excess Ran-GTP in germ cells did not result in severe germline defects (Figure 4D-E, Table 1). The average number of GSCs per germarium was slightly reduced in *nos-Gal4>Ran^Q69L^* ovarioles (Table 1) and a few egg chambers had an incorrect number of cells (Figure 4E), but defects were not as severe or penetrant as either the *Ran* RNAi or Ran-GDP-locked mutants. Since we did not observe a clear Ran intracellular concentration gradient or accumulation of Ran at mitotic spindles, these results suggest that Ran-GTP levels are already in excess in mitotically-active germ cells, perhaps to facilitate timely mitotic divisions.

### Tnpo-SR and cdm, but not Ran-GTP, are required for fusome biogenesis and maturation

The fusome extends through ring canals during incomplete cytokinesis, forming an extensive tubular network whose structure changes over the course of the mitotic divisions (Lighthouse et al., 2008; Röper, 2007; Snapp et al., 2004). The cytoskeletal core is prominent in GSC and cystoblast fusomes, but is replaced with endoplasmic reticulum-like vesicles as cysts exit the germarium (Figure 5A) (Röper, 2007). Loss of *Hts*, a core fusome cytoskeletal protein, alters the number of germ cells per cyst (Yue and Spradling, 1992). Since knock-down of *Ran* or the β-importins produced similar phenotypes, we hypothesized that abnormal fusome morphogenesis might abrogate cyst formation. Using Hts immunostaining as an indicator of fusome morphology, we observed that when *cdm* (Figure 5B) or *Tnpo-SR* (Figure 5C and Supp. Figure S3) were depleted, germ cells formed long, thin fusomes with irregular branching, particularly in cysts in the middle to posterior third of germaria. Fusomes in *Tnpo-SR-* and *cdm*-depleted germ cells often had thread-like sections separated by thicker sections, consistent with weak points in the cytoskeletal structure (Figure 5B-C). Irregular fusomes were also observed in *Tnpo-SR-depleted* GSC/cystoblast pairs, where new fusome material builds from the cystoblast to the GSC (compare Figure 5E-F) (de Cuevas and Spradling, 1998). Importantly, we did not observe fusomes connecting GSCs to other cyst cells in stem-cysts, as was reported for germ cells in which the chromosomal passenger complex, which regulates abscission timing in GSCs, is abrogated (Mathieu and Huynh, 2017). We also did not observe GSCs or cysts with asynchronous mitoses (compare Figure 5G-H), as we would predict if fusomes lost complete connectivity between cells. We conclude that *Tnpo-SR* and *cdm* promote fusome maintenance or remodeling.

**Figure 5.**
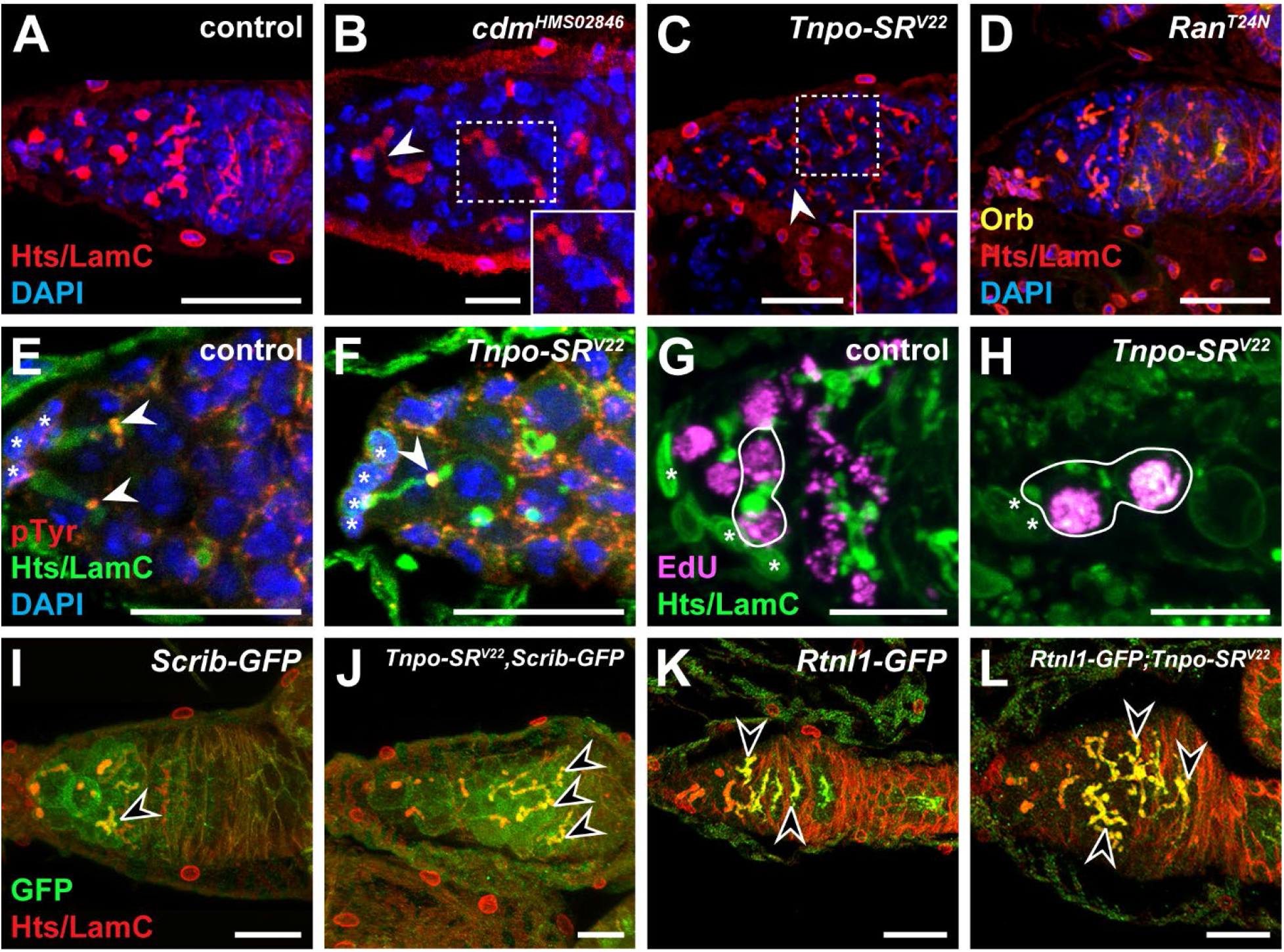
*Tnpo-SR* and *cdm* are required for fusome integrity and maturation. (A-D) Driver control (A), *cdm^HMS02846^* (B), *Tnpo-SR^V22^* (C), and *Ran^T24N^* (D) germaria labeled with anti-Hts (red), anti-LamC (red), and DAPI (blue). Arrowheads indicate points of the fusome that are thinner than normal. (E-F) Driver control (E) and *nos-Gal4>Tnpo-SR^V22^* (F) GSC/CB pairs labeled with anti-pTyrosine (red), anti-Hts (green), anti-LamC (green), and DAPI (blue). Arrowheads indicate pTyr+ plug that forms between the GSC/CB pair prior to abscission. (G-H) Control (G) and *nos-Gal4>Tnpo-SR^V22^* (H) GSC/CB pairs labeled with anti-EdU (magenta), anti-Hts (green) and anti-LamC (green). (I-J) *Scrib-GFP* control (I) and *Tnpo-SR^V22^;Scrib-GFP* (J) germaria labeled with anti-GFP (green), anti-Hts (red), and anti-LamC (red). (K-L) *Rtnl1-GFP* control (K) and *Tnpo-SR^V22^,Rtnl1-GFP* (L) germaria labeled with anti-GFP (green), anti-Hts (red), and anti-LamC (red). Arrowheads indicate highest level of expression of the Scrib (I-J) and Rtnl1 (K-L). All images are maximum intensity projections. Scale bars, 10 μm.

To further investigate how the importins regulate fusome morphogenesis, we recombined the *Tnpo-SR^V22^* RNAi transgene with two fluorescently tagged fusome-resident protein reporters. *Scribble (scrib)* encodes a scaffolding protein that regulates vesicle trafficking and is expressed at the cell cortex and in immature fusomes of GSCs and dividing cysts (Figure 5I) (Lighthouse et al., 2008). By the third mitotic division, *Scrib-GFP* is largely absent from wild-type fusomes and does not co-localize with Hts in 16-cell cysts. In *Tnpo-SR*-depleted germ cells, *Scrib-GFP* was prominently expressed in the long, thin fusomes of germ cells in the posterior of germaria (Figure 5J). Conversely, Reticulon-like1 (Rtnl1) is an endoplasmic reticulum-resident protein that is expressed in low levels in immature fusomes (Figure 5K) (Röper, 2007). *Rtnl1-GFP* becomes enriched in mature fusomes coincident with cyst division and inversely correlated with Hts expression, reaching peak concentration in fusomes after the final mitotic division (Figure 5K). When *Tnpo-SR* is depleted, *Rtnl1-GFP* and Hts co-localize in fusomes throughout the posterior of the germarium (Figure 5L). These data suggest that in the absence of β-importins, cytoskeletal components of the fusome fail to degrade properly, maintaining the fusome in an immature state that is weakly connected between cyst cells. Intriguingly, fusome shape and branching in cysts over-expressing the dominant negative *Ran^T24N^* mutant (Figure 5D) were indistinguishable from wild-type cysts, suggesting that Cdm and Tnpo-SR regulate fusome morphogenesis independently of Ran-GTP.

### Tnpo-SR and cdm promote cyst formation by suppressing cytokinesis

Knock-down of *Tnpo-SR* or *cdm* produced egg chambers variable numbers of germ cells; however, it remained unclear to us exactly how such egg chambers could arise. Key observations came by closely examining the morphology of *Tnpo-SR* mutant cysts surrounded by wild-type cysts in mutant mosaic germaria. While knock-down of *Tnpo-SR* by RNAi led to >16 germ cells per egg chamber, we only rarely found examples of *Tnpo-SR^KG04870^* or *Tnpo-SR^LL05552^* mutant egg chambers with more than 16 mutant germ cells. We did find, however, compound egg chambers in which a *Tnpo-SR* mutant cyst was encapsulated together with wild-type cysts (Supp. Figure S3A). We occasionally identified mitotically-dividing *Tnpo-SR^KG04870^* and *Tnpo-SR^LL05552^* mutant germ cells in mosaic egg chambers outside of the germarium, which never occurs in wild-type germ cells (Supp. Figure S3B). We also frequently found clusters of single *Tnpo-SR* mutant germ cells at the posterior of germaria (Supp. Figure S3C,E-E’’’), suggesting that some cyst defects in *Tnpo-SR* mutant and RNAi-knock-down ovarioles could arise by cyst fragmentation. In the *Tnpo-SR* RNAi model, where all germ cells lack appropriate Tnpo-SR levels, fragmented cysts would likely be encapsulated along with adjacent cysts, thus yielding irregular (i.e., not 2^n^) numbers of germ cells per cyst. In support of this idea, we observed examples of mutant mosaic germaria in which *Tnpo-SR^KG04870^* mutant cells with dot-like fusomes were adjacent to one another with fusomes aligned along the same plane, as if they were connected (Supp. Figure S3D). Immunostaining with anti-pTyr antibodies to visualize ring canals highlighted *Tnpo-SR^KG04870^* mutant cysts where ring canal components accumulated on adjacent cyst cells, separated by a very thin line of GFP-positive cytoplasm (Figure 6A and Supp. Figure S3E), suggesting that cells have physically separated from each other after clone induction. Moreover, *Tnpo-SR^KG04870^* mutant germ cells failed to accumulate proper levels of the cell adhesion protein E-cadherin between germ cells, as occurs in adjacent wild-type 16-cell cysts following the final mitotic division and prior to follicle cell encapsulation (Figure 6B) (Fichelson et al., 2010).

**Figure 6.**
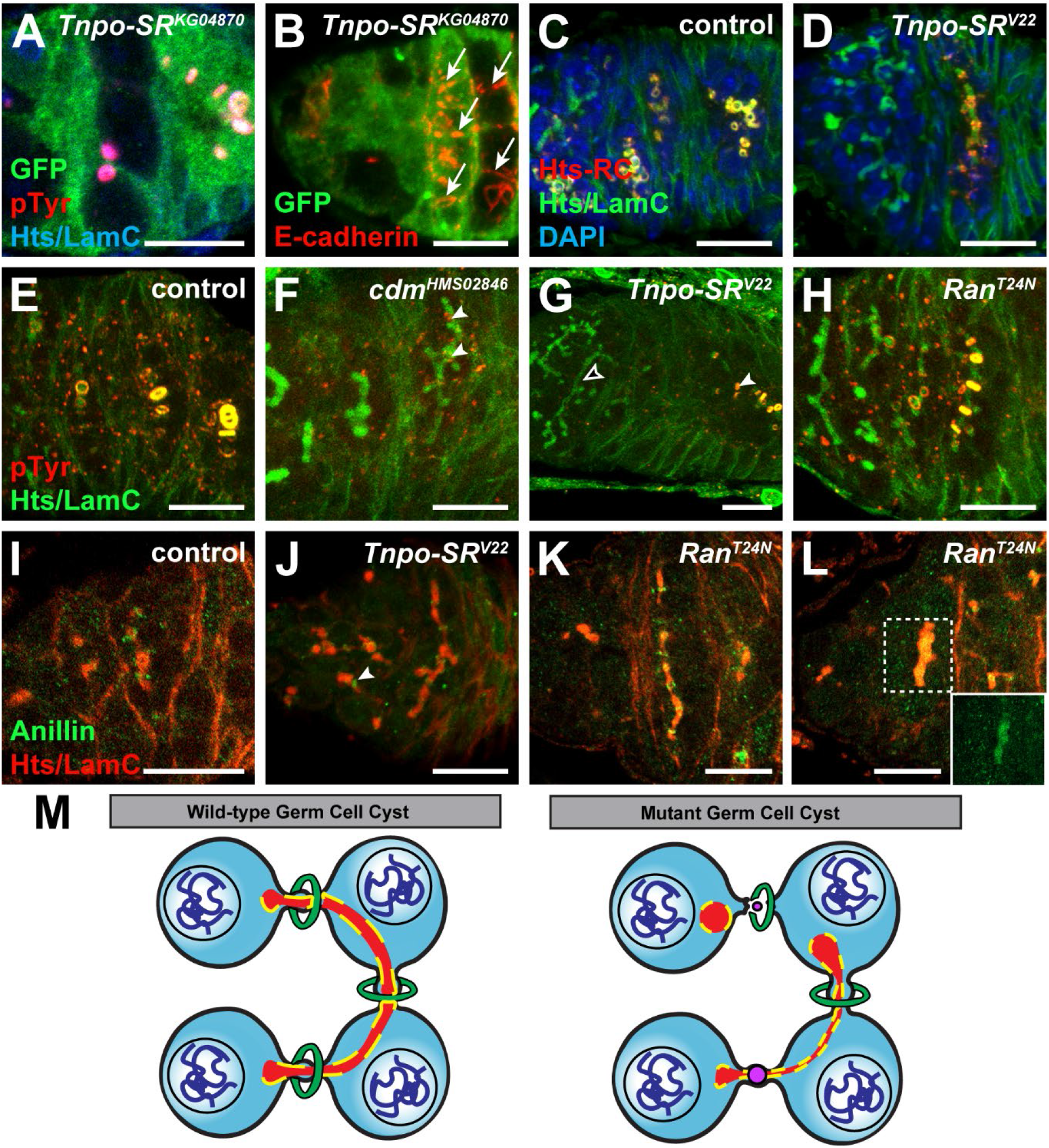
Depletion of *Tnpo-SR* and *cdm*, but not Ran-GTP, delays ring canal protein deposition and fails to suppress cytokinesis in dividing cysts. (A) *Tnpo-SR^KG04870^* mutant mosaic germarium labeled with anti-GFP (green; wild-type cells), anti-pTyrosine (pTyr, red), anti-Hts (blue), and anti-LamC (blue). (B) Maximum intensity projection of a *Tnpo-SR^KG04870^* mutant mosaic germarium labeled with anti-GFP (green; wild-type cells) and anti-E-cadherin (red). Arrows indicate E-cadherin connections. (C-D) Control (C) and *nos-Gal4>Tnpo-SR^V22^* (D) germaria labeled with Hts-RC (red), anti-Hts (green), anti-LamC (green), and DAPI (blue). (E-H) Maximum intensity projections of driver control (E) and RNAi-depleted *cdm* (F), *Tnpo-SR* (G), and *Ran* (H) cysts labeled with anti-pTyr (red), anti-Hts (green), and anti-LamC (green). Arrowheads indicate ring canals; open arrowheads indicate a cyst lacking ring canals. (I-L) Control (I), *nos-Gal4>Tnpo-SR^V22^* (J) and *nos-Gal4>Ran^T24N^* mutant (K-L) germarium labeled with anti-anillin (green) anti-Hts (red), and anti-LamC (red). Arrowheads indicate closed ring canals. Scale bars, 10 μm. (M) Summary of fusome and ring canal phenotypes in *Tnpo-SR*- and *cdm*-depleted germ cell cysts.

These data raised the possibility that in the absence of *Tnpo-SR*, cysts fail to stay connected by intercellular bridges, and instead break apart at unstable ring canals. Initially, this seemed unlikely, as our analyses of *Ran*, *cdm*, and *Tnpo-SR* RNAi and *Ran* dominant-negative germ cells focused on egg chambers outside of the germarium, where ring canals containing Hts-RC (a constituent protein of mature ring canals) were easily detected (Figure 1E-G). Moreover, the number of ring canals scaled to the number of germ cells, suggesting that ring canals formed at successive divisions. Spatiotemporal comparison of ring canal formation, however, demonstrated that while Hts-RC is expressed in germ cells from *nos-Gal4>Tnpo-SR^V22^* ovarioles, its accumulation at ring canals is drastically delayed, such that protein levels are much lower than in age-matched wild-type cysts (compare Figure 6C and D). Similarly, germ cells depleted of *cdm* (Figure 6F) or *Tnpo-SR* (Figure 6G), failed to accumulate pTyr-reactive material (one of the first indicators of ring canal maturation) to wild-type levels in 16-cell cysts. In contrast, ring canal maturation proceeded normally in *Ran^T24N^* mutant cysts (Figure 6H), supporting the model that Tnpo-SR and Cdm, but not Ran-GTP, stabilize germ cell intercellular bridges, maintaining proper cyst formation (Figure 6M).

We then performed immunostaining for Anillin, an actin/myosin-binding protein known to be a β-importin cargo (Beaudet et al., 2017; Beaudet et al., 2020; Chen et al., 2015; Piekny and Maddox, 2010; Silverman-Gavrila et al., 2008). Anillin localization in ring canals begins at the third mitotic division (Figure 6I) (Goldbach et al., 2010; Haglund et al., 2010). In *Tnpo-SR-* depleted germ cells, we observed Anillin deposition within the intercellular bridge, rather than on surrounding rings (Figure 6J). In contrast, Anillin localizes properly to the ring canals in *Ran^T24N^* mutants (Figure 6K) and occasionally co-localized with the fusome (Figure 6L), supporting the model that Tnpo-SR suppresses cytokinesis in dividing cysts independently of Ran-GTP. Intriguingly, over-expression of a GFP-tagged Anillin in which a nuclear localization signal has been mutated did not alter Anillin localization in abscising GSC/CB pairs (Supp. Figure 4A-B) or at ring canals (Supp. Figure 4C-D), and did not change the number of germ cells per cyst. Although we cannot exclude the possibility that Tnpo-SR may bind Anillin at another protein domain, we speculate that Tnpo-SR indirectly regulates Anillin by binding other cargo proteins that impact Anillin localization.

### Ectopic CycA accumulates in Tnpo-SR- and cdm-depleted germ cells

The cell cycle is tightly regulated to coordinate cyst formation. For example, loss of function of the mitotic cyclin CycB produces cysts with eight cells, while over-expression of CycA or CycE produces cysts with 32 cells (Lilly et al., 2000; Mathieu et al., 2013; Morris et al., 2005; Ohlmeyer and Schüpbach, 2003). CycA and CycB accumulate at the fusome and phosphorylation of CycB by AuroraB Kinase promotes abscission between GSC/cystoblast pairs (Lilly et al., 2000; Mathieu et al., 2013). Given the phenotypic parallels, we hypothesized that Tnpo-SR and Cdm might regulate cyclin dynamics in dividing germ cells. We confirmed that CycA accumulates at the fusome in ~10% of mitotically dividing cysts, but never associates with the fusome of 16-cell cysts in wild-type tissue (Figure 7A,F). *Ran* depletion in *nos-Gal4>Ran^GL01341^* germaria did not alter CycA kinetics (Figure 7B,F). In contrast, nearly 40% of mitotically-dividing cysts in *Tnpo-SR* and *cdm*-depleted germ cells expressed CycA; moreover, CycA expression was observed in the fusomes of 16-cell cysts at an equivalent rate as dividing cysts, suggesting that germ cells remain mitotic (Figure 7C-D,F).

**Figure 7.**
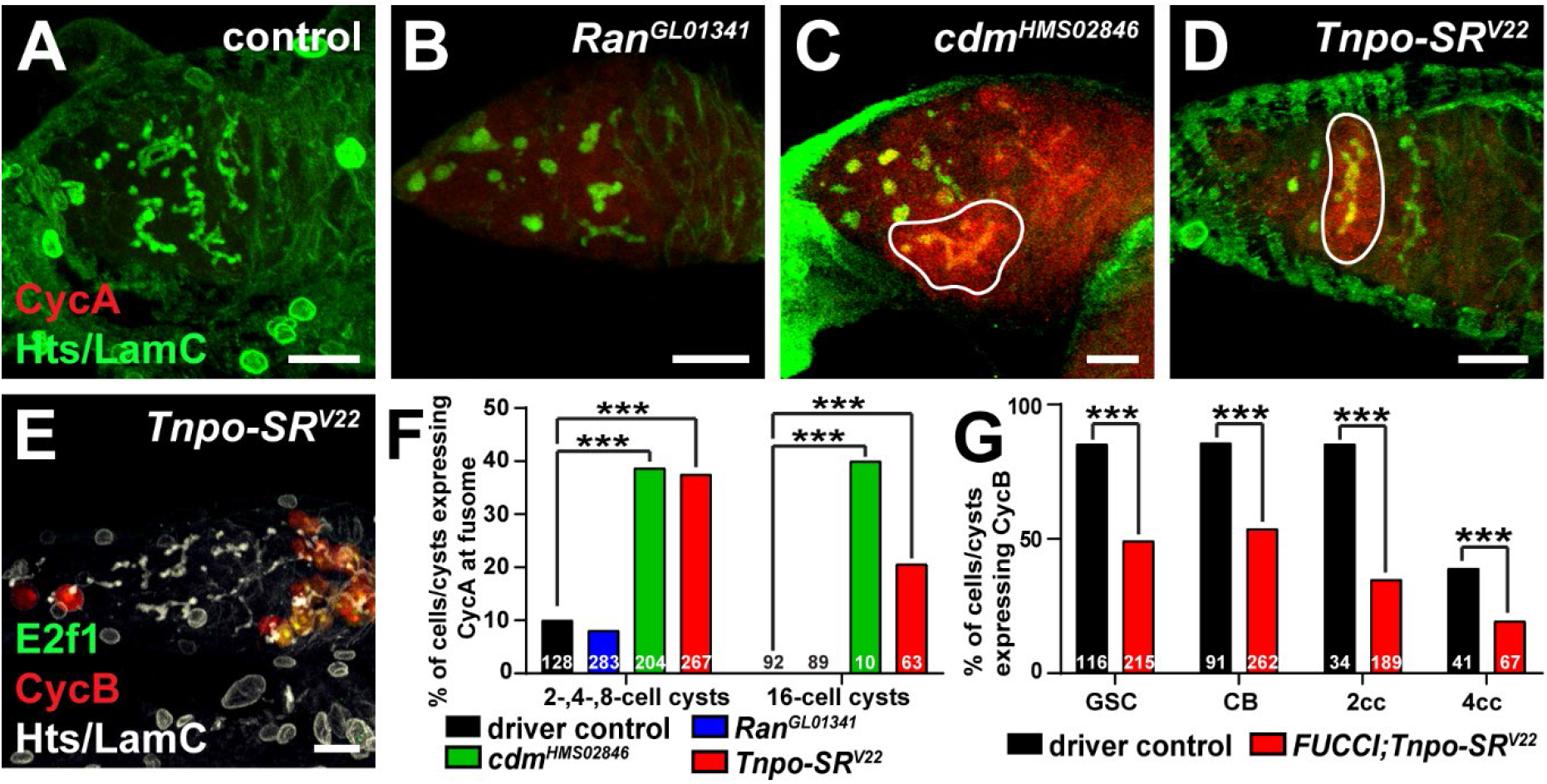
CycA is ectopically expressed in c*dm* and *Tnpo-SR* cysts. (A-D) Driver control (A), *Ran^GL01341^* (B), *cdm^HMS02846^* (C), and *Tnpo-SR^V22^* (D) RNAi knock-down germaria labeled with anti-CycA (red), anti-Hts (green), and anti-LamC (green). (E) *nos-Gal4>FUCCI;Tnpo-SR^V22^* labeled with anti-GFP (green; E2f1), anti-dsRed (red; CycB), anti-Hts (white), and anti-LamC (white). All images are maximum intensity projections. Scale bars, 10 μm. (F) Percentage of cells/cysts containing fusome-localized CycA. (G) Percentage of cells/cysts containing *RFP::CycB* in *nos-Gal4* controls and *Tnpo-SR^V22^* knock-down germaria. The number of cells/cysts analyzed is shown above bars. \*\*\**p* < 0.001, Student’s Chi-squared test.

As the highest peak of CycB expression occurs following CycA degradation (Morris et al., 2005), we then asked whether CycB regulation was likewise impacted by loss of the β-importins. We utilized the *Drosophila* Fluorescence Ubiquitin-based Cell Cycle Indicator system (Fly-FUCCI) (Zielke et al., 2014) to ask whether CycB expression was altered in the absence of *Tnpo-SR*. Importantly, we used a version of the *UASp-CycB-RFP* transgene that includes a canonical nuclear localization sequence (Hinnant et al., 2017; Zielke et al., 2014). When *Tnpo-SR* was depleted, the *UASp-CycB-RFP* localized to germ cell nuclei; however, significantly fewer mitotically dividing cysts expressed CycB as compared to wild-type controls (Figure 7E,G). Taken together, these data suggest that *Tnpo-SR*, and likely *cdm*, promote proper cyclin dynamics in germ cells, maintaining cysts in the mitotic expansion phase and facilitating coordination between cyst division and fusome morphogenesis.

## DISCUSSION

During germline cyst formation, cell cycle transitions are precisely controlled to promote mitotic divisions and block cellular abscission. Our study adds new details to our understanding of the molecular regulation of germ cell mitotic expansion, identifying previously undescribed roles for β-importins and the small GTPase Ran in mitotic cell cycle exit during cyst formation. Remarkably, we do not detect a clear cytoplasmic-to-nuclear gradient of Ran protein localization in germ cell nuclei. Nor do we find defects in mitotic spindle assembly, cell size, or DNA condensation in the absence of β-importins or activated Ran. Instead, our data suggest that importins and Ran-GTP promote mitotic exit by distinct, yet complimentary mechanisms. We propose the model that Ran-GTP drives germ cell mitoses by mobilizing cell cycle factors, functioning together with the β-importins Tnpo-SR and Cdm to facilitate transport of cell cycle factors from the fusome to other parts of the cell. We propose that independent of Ran, Tnpo-SR and Cdm sequester cell cycle regulators like CycA and Anillin, titrating the levels of protein accessible to Ran-GTP, counterbalancing mitotic progression. Thus, in the absence of Tnpo-SR or Cdm, CycA levels accumulate, causing germ cells to continue in a proliferative state and impairing fusome stability, ring canal maturation, and oocyte terminal differentiation.

### Importins may exhibit non-redundant functions in the Drosophila germline

Although several α-importins have been implicated in oogenesis prior to our study (Geles and Adam, 2001; Mason et al., 2002; Máthé et al., 2000), our study is the first to identify specific roles for β-importins in mitotic expansion. Our data generally support the hypothesis that specific importins regulate distinct aspects of germ cell development (Hogarth et al., 2005; Loveland et al., 2015; Mihalas et al., 2015). We cannot exclude the possibility, however, that RNAi was not sufficient to completely inhibit importin function. For example, Ketel localizes at the nuclear envelope in early embryos and dividing germ cells (data not shown) (Lippai et al., 2000; Tirián et al., 2000; Trieselmann and Wilde, 2002; Villányi et al., 2008), yet we did not observe germ cell mitotic defects in *Ketel*-depleted germ cells. Future studies will need to employ other loss-of-function approaches to test the roles of Ketel and other β-importins more thoroughly.

The repertoire of cargo carried by a β-importin determines its role in biological processes and its redundancy with other importins (Kimura and Imamoto, 2014). Although the full extent of protein-protein interactions mediated by Tnpo-SR remains unknown, structural studies indicate that TNPO3 binds proteins involved in chromatin organization, mRNA and protein modification, cell cycle control, and cell-cell communication (Kimura et al., 2017; Maertens et al., 2014). Tnpo-SR is also likely to bind a broad range of cargo proteins, including SR-rich RNA binding proteins. Indeed, Tnpo-SR physically interacts with several proteins essential for mRNA splicing and transport (Allemand et al., 2002). Given that SR-rich proteins facilitate diverse RNA processing events, there are many potential Tnpo-SR cargo proteins that could influence cyst formation and GSC self-renewal, which are heavily influenced by translational control (Bradley et al., 2015; Teixeira and Lehmann, 2019). Our future studies will seek to identify Tnpo-SR and Cdm-interacting proteins, which should provide additional mechanistic details regarding how importins regulate germ cell mitotic expansion.

### The number of cyst divisions is regulated by Ran-GTP and β-importins

Precise spatiotemporal control over cell cycle regulators underlie the transition to the oocyte fate. In support of this, the mitotic cyclin CycA is clearly critical to support differentiation. Failure to degrade CycA leads to self-renewal failure in GSCs, and extra mitotic divisions and loss of oocyte identity in cyst cells (Chen et al., 2009; Ji et al., 2017; Lilly et al., 2000; Morris et al., 2005; Ohlmeyer and Schüpbach, 2003; Sugimura and Lilly, 2006). Moreover, the deubiquitinase encoded by *bag of marbles (bam)*, one of the few known differentiation factors in *Drosophila* germ cells, is necessary for the transition from GSC to cystoblast and for cystoblast divisions, at least in part because it stabilizes CycA expression in dividing cysts (Ji et al., 2017). CycA/Cdk1 activity is very high in early cyst divisions but decrease as cysts approach the terminal mitotic division (Hinnant et al., 2017; Lilly et al., 2000). Since cystoblasts divide exactly four times, prior studies posited the model that cystoblasts autonomously limit the number of divisions through a molecular counting mechanism that involves Bam, CycA, and the fusome (Hinnant et al., 2020; Huynh, 2006; Ji et al., 2017; King, 1970; Lilly et al., 2000; McKearin, 1997). While a counting mechanism would limit the number of mitotic divisions, it must also promote modification of contractile ring proteins to block cytokinesis, stabilize intercellular bridges into stable ring canals, and maintain the biosynthesis of the fusome (McKearin, 1997).

We suggest that the Ran/β-importin intracellular transport system is integrated into the CycA-dependent cell division counting machinery. Depletion of *Ran*, *cdm*, or *Tnpo-SR* generated egg chambers with abnormal numbers of cells (Table 1), some on the order of 2^n^ reminiscent of cyclin mutants (Lilly et al., 2000; Mathieu et al., 2013; Morris et al., 2005; Ohlmeyer and Schüpbach, 2003), and others deviating from this pattern, more similar to mutants which disrupt intercellular bridge formation and fusome biogenesis (de Cuevas et al., 1996; Lin et al., 1994; Yue and Spradling, 1992). Although we cannot establish a clear temporal sequence of events mediated by Ran from our current data, we envision two possible mechanisms of action. We favor the hypothesis that Ran mediates intracellular movement of CycA itself or a factor that promotes CycA degradation. Binding by Tnpo-SR or Cdm could sequester the cell cycle regulator, preventing activation by sterically inhibiting interaction with a kinase or by blocking protein accumulation at the fusome. High levels of Ran-GTP levels would promote release of the Tnpo-SR cargo, freeing it for activation. In the absence of Tnpo-SR or Cdm, where mitotic regulators cannot be delivered to Ran, CycA levels instead accumulate at the fusome and in the cytoplasm, promoting continued mitotic division. Alternatively, Ran-GTP could regulate contractile protein localization and cell polarization, as has been demonstrated for human Ran in mammalian somatic cells (Beaudet et al., 2017; Silverman-Gavrila et al., 2008).

### Tnpo-SR and Cdm may have Ran-independent roles in promoting cyst development

Given the close association between Ran and β-importins, we hypothesized that depletion of *Ran* by RNAi, or over-expression of dominant negative *Ran^T24N^* mutant would phenocopy β-importin depletion. Yet we observed phenotypes unique to loss of *Tnpo-SR* and *cdm*. In *Tnpo-SR-* and *Cdm-* depleted germ cells, delayed maturation of the fusomes and ring canals appears to create weak connections between cells, resulting in premature abscission or collapsed intercellular bridges (Figure 6M). Interestingly, we see fusomes that are fragile and unbranched (Figure 5E-H), but also germline tumors containing cells with immature fusome morphology resembling that of 2-cell cysts (Figure 2G-H). This may be the result of altered timing or different concentrations of the β-importins in the first division of the cystoblast versus the remaining three, perhaps reflecting differing sensitivity to abscission across the mitotic divisions (Mathieu et al., 2013; McKearin and Ohlstein, 1995). Cyst cell fusomes are composed of endoplasmic reticulum-like vesicles and cytoskeletal elements that originate from remnants of the mitotic spindle midbody (de Cuevas and Spradling, 1998; Koch and King, 1966; Koch et al., 1967; Lin and Spradling, 1995; Mahowald, 1971). This suggests Ran-independent functions of Tnpo-SR and Cdm in mitotically dividing germ cells, perhaps by sequestering negative regulators of fusome maturation or cytokinesis. We speculate that Tnpo-SR and Cdm promote reorganization of the germ cell cytoskeleton after mitotic division, perhaps by shuttling cytoskeletal proteins back to the fusome.

### Ran-GTP and β-importins may function as a conserved molecular timer for the mitotic-to-meiotic transition

The basic steps in cyst formation have been well described at the morphological level, particularly in insects (de Cuevas et al., 1997; Lu et al., 2017; Pepling and Lei, 2018; Yamashita, 2018). Yet how cyst formation is guided molecularly remains an open area for study. Insects provide excellent models, as many exhibit unique mechanisms for achieving cyst formation (Büning, 1994; de Cuevas et al., 1997; Eastin et al., 2020; Kubrakiewicz, 1997). Given the level of conservation in cell cycle control across species, we propose that Ran and β-importins may also play important roles in higher eukaryotes, connecting the intricate programs of cell division, oocyte differentiation, and cell polarization. This may be of particular consequence for mouse and human oocytes, in which cyst breakdown occurs prior to meiotic onset (Pepling and Lei, 2018). Understanding how Ran and importins control cyst formation and breakdown may help elucidate how the oocyte reserve is established in humans, offering new potential avenues for treatment of infertility.

## MATERIALS AND METHODS

### Drosophila strains and husbandry

Flies were maintained at 22°-25°C in standard medium (cornmeal/molasses/yeast/agar) (NutriFly MF; Genesee Scientific). Female progeny were collected one to two days after eclosion and maintained on standard medium for 3, 6, and 10 days at 25°C; flies were fed wet yeast paste for 2-3 days (changed daily) prior to ovary dissection. A complete list of fly lines used in this study are provided in Supp. Table S1.

### Tissue-specific RNA interference (RNAi) generation

All RNAi lines used in this study were carried in *pVALIUM20* or *pVALIUM22* transposons for maximum germline efficiency (Supp. Table S1). *UAS-Tnpo-SR^V22^* was generated as described (www.flyrnai.org/TRiP-HOME.html) (Ni et al., 2011). Primers were designed against the full-length *Tnpo-SR* RNA via the Designer of Small Interfering RNA tool (http://biodev.extra.cea.fr/DSIR/DSIRhtml) using default settings (21 nt siRNA; score threshold 90). A sequence with a score of 94.6 and 0 predicted off-targets was selected for hairpin generation (passenger strand, CGATCCCGTTTACTGGATAGA; guide strand, TCTATCCAGTAAACGGGATC). Primers were designed according to TRiP recommendations, annealed to form double-stranded oligos, and ligated into EcoRI-HF/NheI-HF-digested *pVALIUM22*. Transgenic flies *(UAS-Tnpo-SR^V22^)* were generated by phiC31 site-specific integrase into the *attP2* site on the third chromosome.

For RNAi experiments, germline knock-down was facilitated by expressing the germline-specific *nos-GAL4::VP16-nos.UTR* (referred to throughout as *nos-Gal4*) (Rørth, 1998; Van Doren et al., 1998). Driver expression was confirmed using *w*; nos-Gal4, UASp-tubGFP* (BDSC #7253) (Grieder et al., 2000). Females carrying *nos-Gal4* alone were used as controls.

### Drosophila genetic mosaic generation

For genetic mosaic analyses using *flippase (FLP)/FLP recognition target (FRT)*, we obtained the following mutant alleles of *FRT*-containing chromosome arms: *Tnpo-SR^KG04870^*[Kyoto *Drosophila* Stock Center (KDSC) #111581] (Bellen et al., 2004) and *Tnpo-SR^LL05552^*(KDSC #141628) (Schuldiner et al., 2008). Other genetic tools are described in FlyBase version FB2020_05 (Thurmond et al., 2019). Genetic mosaics were generated by *FLP*/*FRT*-mediated recombination in 2-3-day old females carrying a mutant allele in trans to a wildtype allele (linked to a *Ubi-GFP* marker) on homologous *FRT* arms, and a *hs-FLP* transgene, as described (Laws and Drummond-Barbosa, 2015). Flies were heat shocked at 37°C two times per day for 3 days and incubated at 25°C for 4, 8, or 12 days and supplemented with wet yeast paste on the last 2 days prior to dissection. Wild-type alleles were used for the generation of control (mock) mosaics.

### Germ cell analyses

GSCs were identified based on the juxtaposition of their fusomes to the junction with adjacent cap cells (de Cuevas and Spradling, 1998). In *FLP/FRT* mutants, GSC loss was measured two ways. First, as the percentage of total mosaic germaria showing evidence of recent stem cell loss; namely, the presence of GFP-negative daughters (cystoblasts/cysts generated from an original GFP-negative stem cell) in the absence of the GFP-negative mother stem cell (Laws and Drummond-Barbosa, 2015). Second, as the percentage of total germaria analyzed that had at least one GFP-negative GSC. All results were subjected to Chi-Square analyses using Microsoft Excel. In RNAi progeny, GSC loss was measured as the average number of GSCs present over three timepoints. Results were subjected to Student’s two-tailed T-test comparing driver controls versus *driver>UAS-RNAi* experimental groups at each timepoint individually using Microsoft Excel.

For analyses involving the egg chambers outside of the germarium, data was collected from the first three egg chambers posteriorly located from the germarium. Oocytes were identified by the presence of anti-Orb antibodies. The number of cells and ring canals per egg chamber were calculated based on visualization of ring canals with anti-pTyr and/or anti-Hts-RC antibodies and cell nuclei with DAPI. Results were subjected to Student’s two-tailed T-test comparing driver controls versus *driver>UAS-RNAi* experimental groups at each timepoint individually using Microsoft Excel and Prism (GraphPad).

### Immunofluorescence and microscopy

Ovaries were prepared for immunofluorescence microscopy as described (Grieder et al., 2000; Hinnant et al., 2017). In the standard protocol, ovaries were dissected and teased apart in Grace’s medium without additives (Caisson Labs) and fixed in 5.3% formaldehyde in Grace’s medium for 13 minutes at room temperature. They were then washed extensively in phosphate-buffered saline (PBS, pH 7.4; Fisher) with 0.1% Triton X-100, and blocked for three hours in blocking solution [5% bovine serum albumin (Sigma), 5% normal goat serum (MP Biomedicals), and 0.1% Triton X-100 in PBS] at room temperature. To detect cells in S phase, dissected ovaries were incubated for one hour at room temperature in Grace's media containing 10 μM 5-ethynyl-2′-deoxyuridine (EdU; Life Technologies). The following primary antibodies were diluted in blocking solution and used overnight at 4°C: chicken anti-GFP (#13970, Abcam; 1:2000), rabbit anti-dsRed (#632496, Clontech; 1:500), rabbit anti-phosphoHistone H3 (pHH3; #06-570, Millipore; 1:200), rat anti-E-Cadherin (DCAD2, Developmental Studies Hybridoma Bank (DSHB); 1:20), mouse anti-Orb (4H8 and 6H4; DSHB; 1:500), mouse anti-BicD (1B11 and 4C2; DSHB; 1:10), mouse anti-p-Tyrosine (Millipore; 1:100), mouse anti-Hts-RC (DSHB; 1:10), mouse anti-CycA (A12; DSHB; 1:50), rabbit anti-C(3)G (a gift from M. Lilly; 1:3000), rat anti-HA (3F10; Sigma; 1:25), mouse anti-α-Tubulin-AF555 (DM1A-AF555; Millipore; 1:100), and rabbit anti-Anillin (a gift from J. Brill; 1:50). Mouse anti-Sxl (m18; DSHB; 1:500) was also used but fixed in a 4% formaldehyde in Grace’s medium. Primary antibodies mouse anti-Hts (1B1, DSHB; 1:10), and mouse anti-Lamin C (LC28.26, DSHB; 1:100) were incubated over two nights at 4°C. All primary antibodies are followed by a two hour incubation at room temperature with AlexaFluor 488-, 568- or 633-conjugated goat species-specific secondary antibodies (Life Technologies; 1:200). EdU was detected (if necessary) using AlexaFluor-594 or −647 via Click-It chemistry, following the manufacturer′s recommendations (Life Technologies).

For stabilizing structures such as mitotic spindles, ovaries were dissected and teased apart in room temperature equilibrated Grace’s medium and fixed for 10 minutes at room temperature using 350μl Grace’s Medium and 200μl of 16% formaldehyde. They were then extensively washed in PBS with 0.1% Triton X-100 and blocked at room temperature for at least 30 minutes in blocking solution (2.5mL NGS, 5mL Triton, 42.5mL 1xPBS). Primary antibody incubations took place overnight at 4°C followed by extensive washing in PBS with 0.1% Triton X-100 and another blocking period of at least 30 minutes at room temperature. Detection of secondary antibodies proceeded as described above.

All ovary samples were stained with 0.5 μg/ml 4’-6-diamidino-2-phenylindole (DAPI; Sigma) in 0.1% Triton X-100 in PBS, and mounted in 90% glycerol mixed with 20% n-propyl gallate (Sigma). Confocal z-stacks (1 μm optical sections) were collected with either the Zeiss LSM700 laser scanning microscope using ZEN Black 2012 software or the Zeiss LSM800 laser scanning microscope using ZEN Blue 2012. Images were analyzed using Zeiss ZEN software, and minimally and equally enhanced via histogram using ZEN and Adobe Photoshop Creative Suite.

## ACKNOWLEDGMENTS

Many thanks to K. McKim, B. Riggs, K. Roper, J.-Q. Ni, M. Lilly, D. Drummond-Barbosa, M. Buszczak, L. Cooley, and J. Brill for their kind gifts of fly stocks and antibodies. We are grateful to the Bloomington, Vienna, and Kyoto *Drosophila* Stock Centers, the Drosophila Genomics Resource Center, the Harvard Transgenic RNAi Project, and the University of Iowa Developmental Studies Hybridoma Bank for making fly lines, antibodies, and plasmids available, and BestGene for *Drosophila* transgenesis and screening. Many thanks to Tianna Van Cura and members of the Ables laboratory for assistance with building tools, helpful discussions, and critical reading of this manuscript.

## DATA AVAILABILITY

Fly strains and plasmids are available upon request. The authors affirm that all data necessary for confirming the conclusions of the article are present within the article and figures.

## SUPPLEMENTAL DATA

**Supp. Figure 1.**
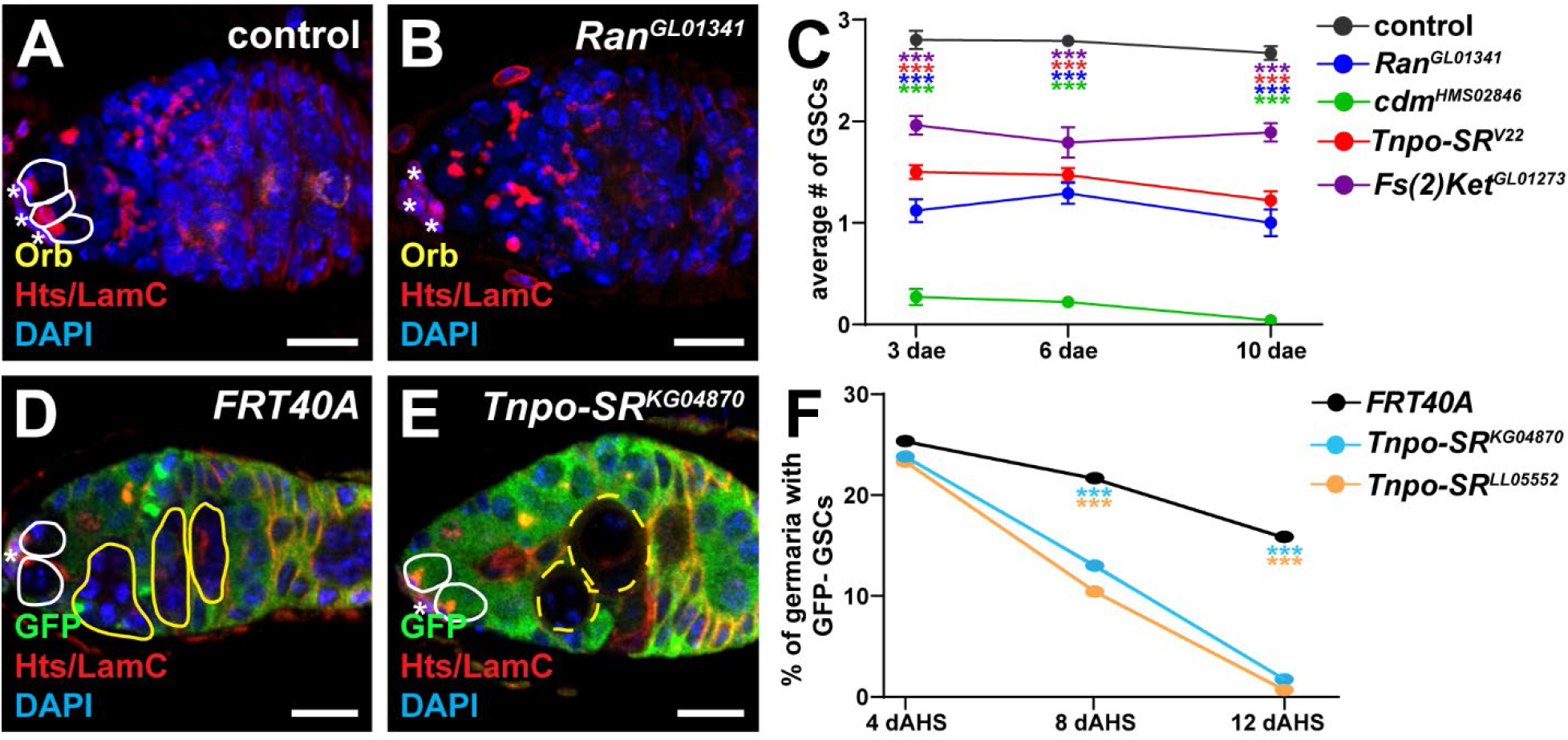
Ran and β-importins are necessary for GSC maintenance. (A-B) *nos-Gal4* control (A) and *nos-Gal4>Ran^GL01341^* (B) germaria labeled with anti-Orb (yellow), anti-Hts (red), anti-LamC (red) and DAPI (blue). Solid white lines demarcate GSCs present in the germarium, asterisks indicate cap cells. (C) Average number of GSCs per germarium in *nos-Gal4* control or *nos-Gal4>UAS-Ran/importin RNAi* ovaries at 3, 6, and 10 days after eclosion (dae). At least 30 germaria were analyzed at each timepoint. (D-E) *FRT40A* control (D) and *Tnpo-SR^KG04870^ FRT40A* mutant (E) mosaic germaria at 8 days after clone induction labeled with anti-GFP (green), anti-Hts (red), anti-LamC (red) and DAPI (blue). Lines demarcate wild-type (solid) or mutant (dashed) GSCs (white) and cystoblast/cysts (yellow). (F) Percentage of total germaria analyzed with a GFP-negative GSC at 4, 8, and 12 days after clone induction. At least 145 germaria were analyzed for each genotype at each timepoint.

**Supp. Figure 2.**
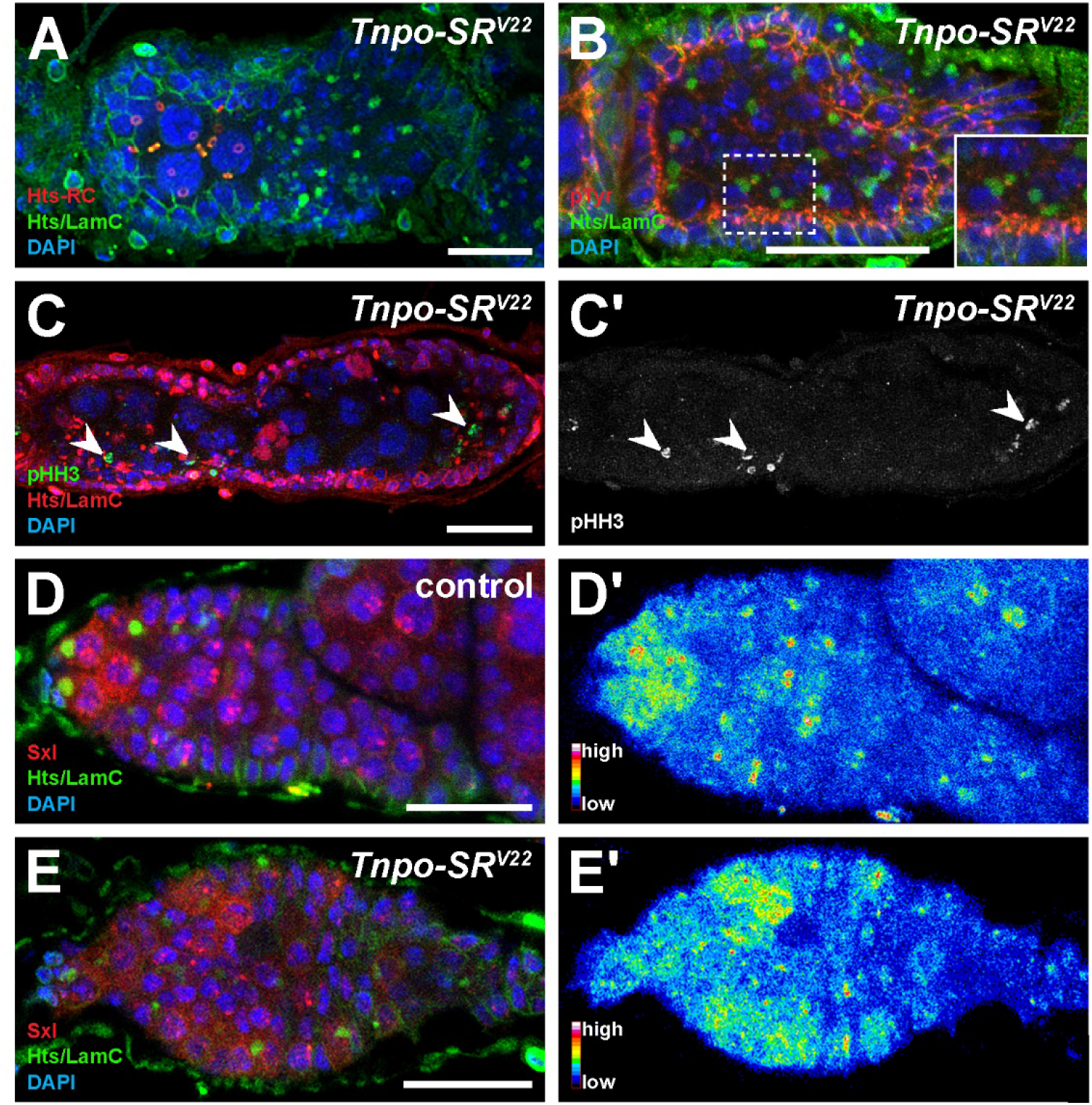
*Tnpo-SR* germline tumors are primarily composed of mitotically-dividing undifferentiated germ cells. (A) *nos-Gal4>Tnpo-SR^V22^* egg chamber labeled with anti-Hts-RC (red), anti-Hts (green), anti-LamC (green), and DAPI (blue). (B) *nos-Gal4>Tnpo-SR^V22^* egg chamber labeled with anti-pTyr (red), anti-Hts (green), anti-LamC (green), and DAPI (blue). (C-C’) *nos-Gal4>Tnpo-SR^V22^* egg chamber labeled with anti-pHH3 (green), anti-Hts (red), anti-LamC (red), and DAPI (blue). Panel C’ depicts pHH3 channel only. Arrowheads indicate pHH3-positive cells within the germline tumor. (D-E) Control (D-D’) and *nos-Gal4>Tnpo-SR^V22^* (E-E’) germaria labeled with anti-Sxl (red), anti-Hts (green), anti-LamC (green), and DAPI (blue). Panels D’ and E’ show heat maps corresponding to panels D and E, where cool colors are low expression and warm colors are high expression. Scale bars, 20 μm.

**Supp. Figure 3.**
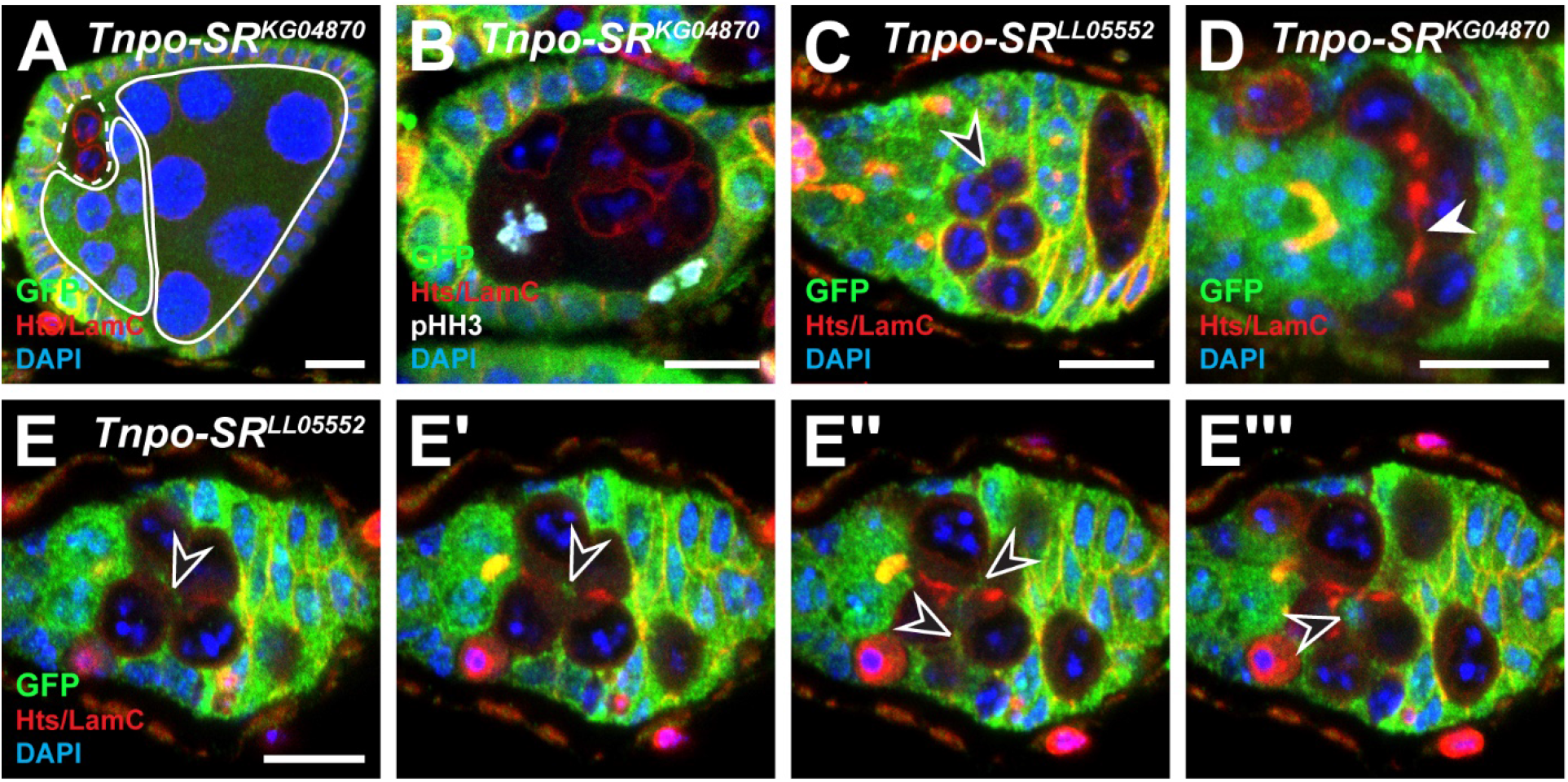
*Tnpo-SR* mutants fail to maintain cyst connectivity and some cells are ectopically mitotically active. (A) *Tnpo-SR^KG04870^* mutant egg chamber labeled with anti-GFP (green), anti-Hts (red), anti-LamC (red), and DAPI (blue). White lines indicate wild-type cysts (solid) and mutant cysts (dashed). (B) *Tnpo-SR^KG04870^* mutant egg chamber labeled with anti-GFP (green), anti-pHH3 (white), anti-Hts (red), anti-LamC (red), and DAPI (blue). (C-E’’’) *Tnpo-SR^LL05552^* (C and E-E’’’) and *Tnpo-SR^KG04870^* (D) mutant germaria labeled with anti-GFP (green), anti-Hts (red), anti-LamC (red), and DAPI (blue). Panels E-E’’’ are adjacent 1 μm optical sections from the same germarium. Arrowhead indicates thin thread of fusome material; open arrowheads follow the GFP+ somatic cell projections invaginating the mutant cysts. (D) is a 2 μm subset maximum intensity projection. Scale bars, 10 μm.

**Supp. Figure 4.**
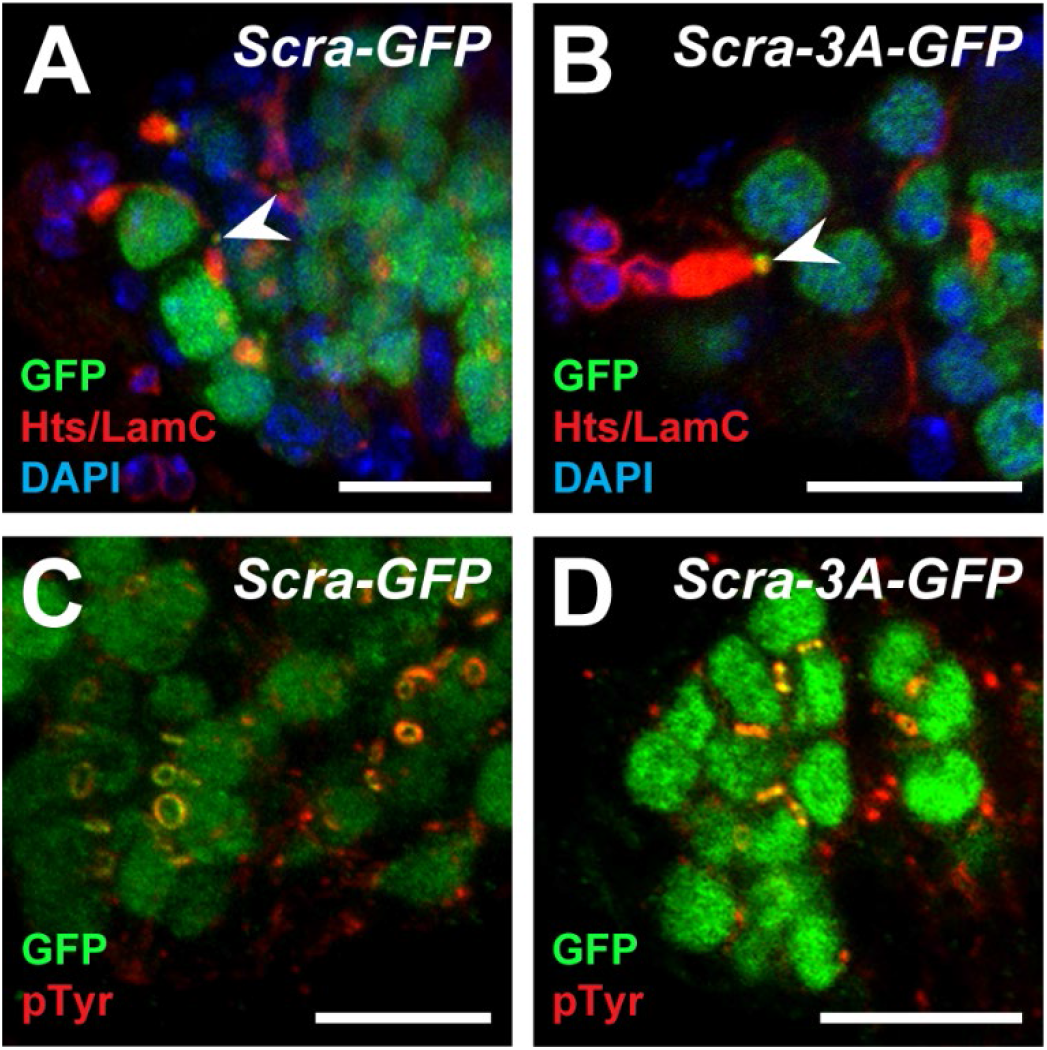
An Anillin-NLS mutant is not efficient in disrupting Anillin localization to nuclei and ring canals. (A-D) Representative controls of *UAS-Scra-GFP* (A and C) and NLS mutated *UAS-Scra-3A-GFP* mutant (B and D) germaria labeled with anti-GFP (green), anti-Hts (red), anti-LamC (red), and DAPI (blue) (A-B) or anti-pTyrosine (red) (C-D). Arrowheads indicate closed ring canals. Scale bar, 10 μm.

**Supp. Table S1.**
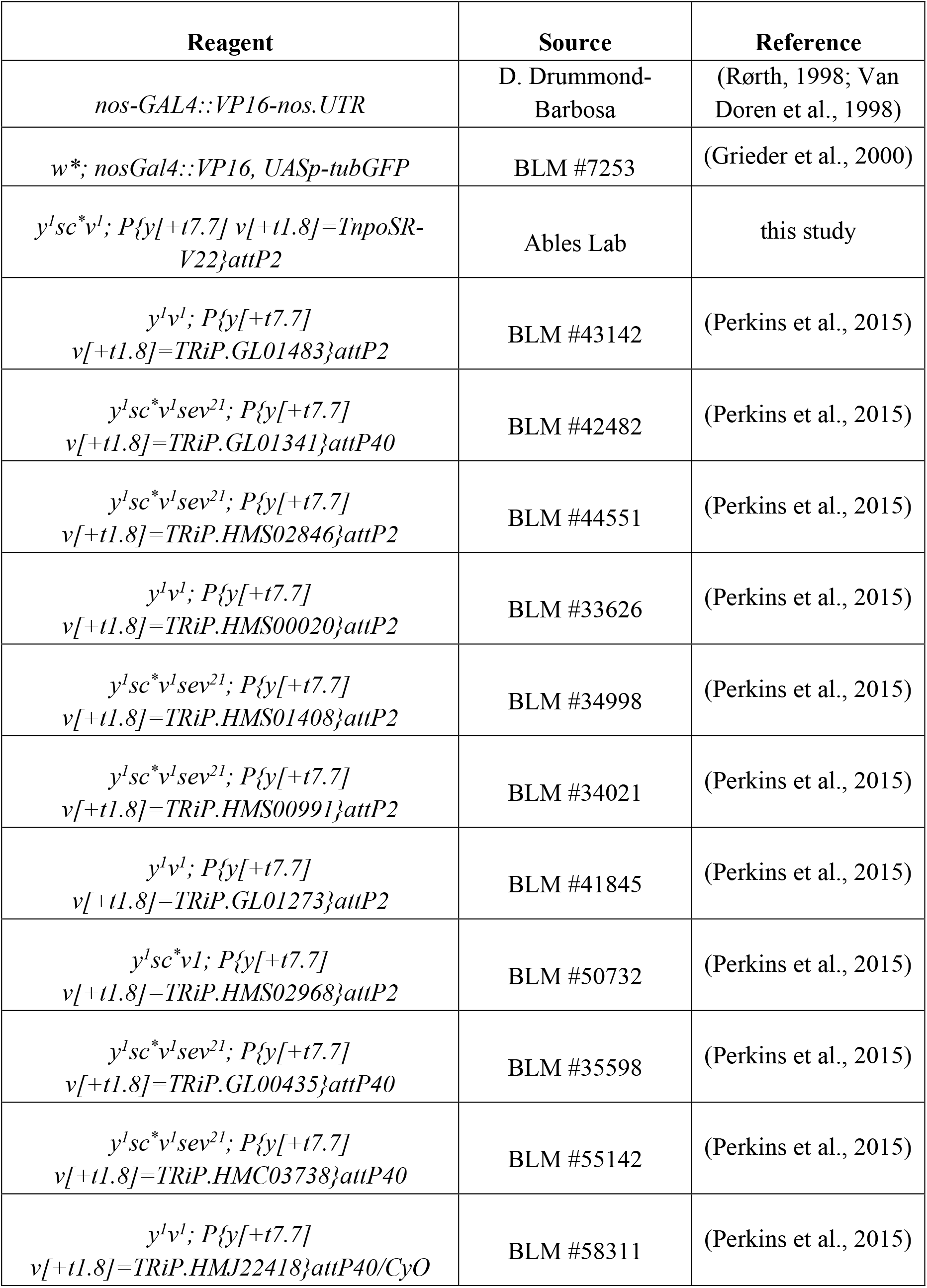
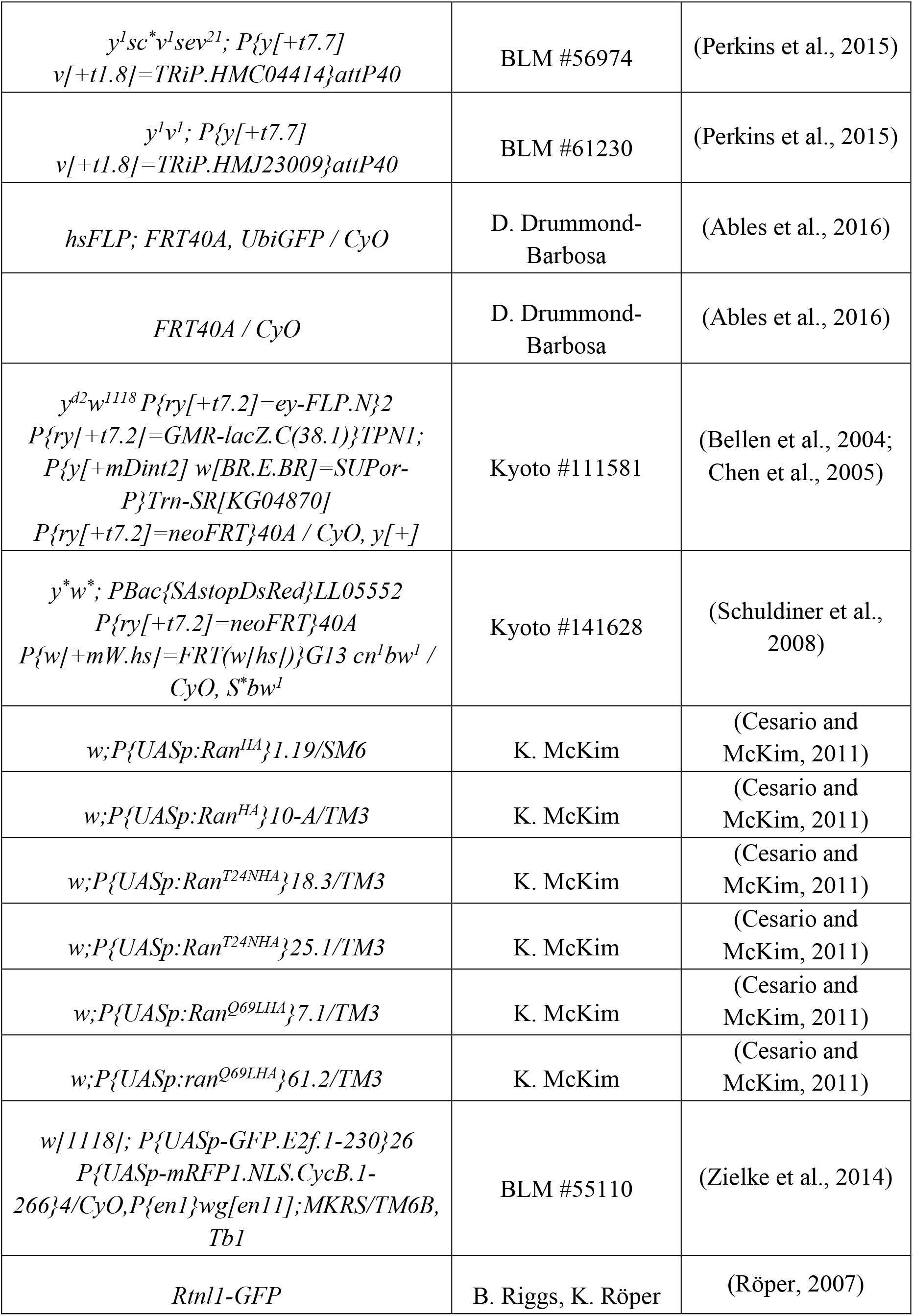
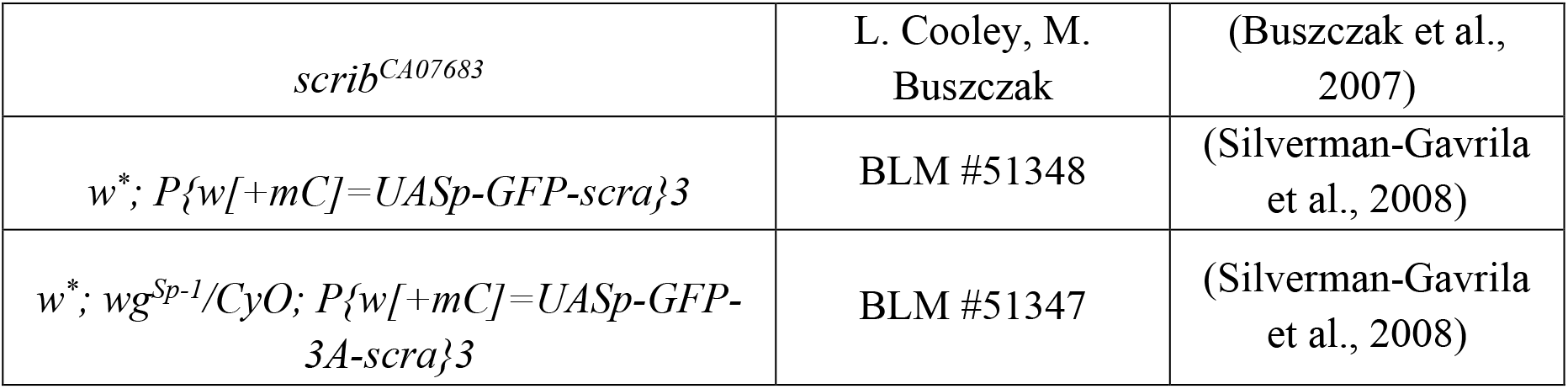
Fly stocks and immunofluorescence reagents used in this study.

